# Habitat fragmentation and food security in crop pollination systems

**DOI:** 10.1101/2020.05.28.120550

**Authors:** Daniel Montoya, Bart Haegeman, Sabrina Gaba, Claire De Mazancourt, Michel Loreau

**Affiliations:** Centre for Biodiversity Theory and Modelling, Theoretical and Experimental Ecology Station, UMR 5321, CNRS, 09200 Moulis, France; USC 1339, Centre d’Etudes Biologiques de Chizé, INRA, 79360 Villiers en Bois, France; Centre d’Etudes Biologiques de Chizé, UMR 7372, CNRS & Université de La Rochelle, 79360 Villiers en Bois, France

## Abstract

Ensuring stable food supplies is recognized as a major challenge for the 21^st^ century, and one of the UN Sustainable Development Goals. Biodiversity-based approaches to food security are increasingly being supported based on the fact that biodiversity can increase and stabilize crop yields. But agricultural systems are often highly fragmented and it is unclear how such fragmentation affects biodiversity and food production, limiting our capacity to manage agricultural landscapes for food security. Here, we develop a model of crop yield dynamics to investigate how fragmentation of natural habitats for agricultural conversion impacts food production, with a focus on crop pollination. Our results show that fragmentation produces spatial and biodiversity-mediated effects that affect the mean and stability of pollination-dependent crops, with strong consequences for food security. The net effects of fragmentation depend on the strength of the spillover of pollinators to crop land and the degree to which crops depend on animal pollination. Our study sheds new light in the food security debate by showing that high and stable yields depend on biodiversity and the spatial structure of agricultural landscapes, and by revealing the ecological mechanisms of food security in crop pollination systems.

## INTRODUCTION

Ensuring stable food supplies for a growing population is one of the UN Sustainable Development Goals^1^, and is a challenge that may require multiple solutions. Biodiversity-based approaches to food security suggest that safeguarding certain levels of species diversity is fundamental to increase yields and guarantee stable yields. This is supported by theory and data reporting positive effects of biodiversity on the mean values of various ecosystem functions and services^2-6^. Additionally, biodiversity can have a stabilizing effect on ecosystem service supply by providing an insurance against environmental fluctuations^7^, which are predicted to intensify under global change^8-11^. Biodiversity insurance effects have been observed in agriculture, where a greater diversity of crops in arable land is associated with increased year-to-year temporal stability of total yields^12^. However, most biodiversity in intensively-managed agricultural landscapes is found in the remaining fragments of (semi)natural habitat not converted into crop land, and the effects of such non-crop biodiversity on both the mean provision and stability of crop production are not clearly understood. This has led to a growing concern over the large-scale conversion of natural habitats into crop land and their effects on biodiversity and food production.

Agricultural systems are often highly fragmented with areas of intensive cultivation interspersed among remnant patches of seminatural habitat. This loss and fragmentation negatively affect biodiversity and many ecosystem functions and services^13-15^. Despite this, fragments of natural habitat continue to supply important services. The spatial coexistence of crops and natural land creates an opportunity for spillover effects^16^, a situation where ecological interactions extend across habitats boundaries and propagate ecological functions. In some cases, fragmentation can increase ecosystem service supply, e.g. if fragmentation of natural habitat for pollinators optimizes interspersion with crop land to maximize crop pollination^17^. But fragmentation can also reduce the provision of ecosystem services if biodiversity decreases significantly in the remnant fragments of natural habitat^13-14^. For most services, however, we do not know how fragmentation affects their provision in fragmented landscapes, and this limits our capacity to manage ecosystem service provision and food security in human-dominated landscapes.

Recent research has revealed strong and non-linear effects of land conversion on agricultural pollination services at multiple spatial scales^18-20^. For example, there is consensus on the hump-shaped relationship between the pollinator-dependent component of crop production and the fraction of remnant natural land within intensive farming systems^20,21^. Research on the stability of crop production shows that yield stability decreases with agricultural intensification and the degree to which crops depend on animal pollination^22-25^. Changes in crop yield stability of animal-pollinated crops are also dependent on the spatial composition and structure of agricultural landscapes, such as the amount of remnant natural land cover^20,26^ and the isolation of crops from natural land^23^. Overall, these studies suggest that improved management of agricultural landscape should increase the amount and stability of production for pollination-dependent crops, and that an understanding of how the spatial pattern of land conversion – i.e. fragmentation – impacts ecosystem services is key to achieve this goal. However, none of these studies have simultaneously combined several ecosystem services, crop yield dynamics at different scales and spatially-explicit landscapes to investigate the effects of land conversion on biodiversity and the provision and stability of ecosystem services in agricultural landscapes.

There is general consensus that increased food production is necessary, but not sufficient, to achieve food security^27^, and that agriculture, especially in the global change context, should also aim at stabilizing crop yields over time^26,28^. Bearing this in mind, we here extend a model of crop yield dynamics into a spatially-explicit landscape to investigate how habitat loss and fragmentation, i.e. the amount and spatial configuration of seminatural habitat, influence the mean provision and stability of several ecosystem services in agricultural landscapes. We focus on crop pollination systems because (i) crop pollination is a key agricultural service that depends on biodiversity, and (ii) worldwide agriculture is shifting towards more pollinator-dependent food production systems^29,30^. Because the way food is produced worldwide threatens the existence of much of the world’s biodiversity that contributes to crop pollination and food security, we explore how changes in biodiversity following land conversion affect the supply of various ecosystem services in fragmented agroecosystems. Specifically, we address two questions: (i) How does the spatial pattern of land conversion, i.e. loss and fragmentation of natural habitat, influence the provision and stability of crop pollination services in agroecosystems? (ii) How does biodiversity in fragmented landscapes influence crop pollination and food security?

## METHODS

### a. Spatial agroecosystem model

We developed a model to investigate the expected biodiversity (i.e. species richness), crop production at the farm level (i.e. crop yield per area) and landscape crop production (i.e. the magnitude and stability of crop pollination and independent crop yield) in agricultural landscapes with varying degrees of fragmentation and for different crop types (i.e. different levels of animal pollination dependence), yielding a total of six ecosystem service components. In what follows, we describe the model dynamics and the land conversion pattern generation. A conceptual representation of our model is provided in Figure 1.

**Figure 1.**
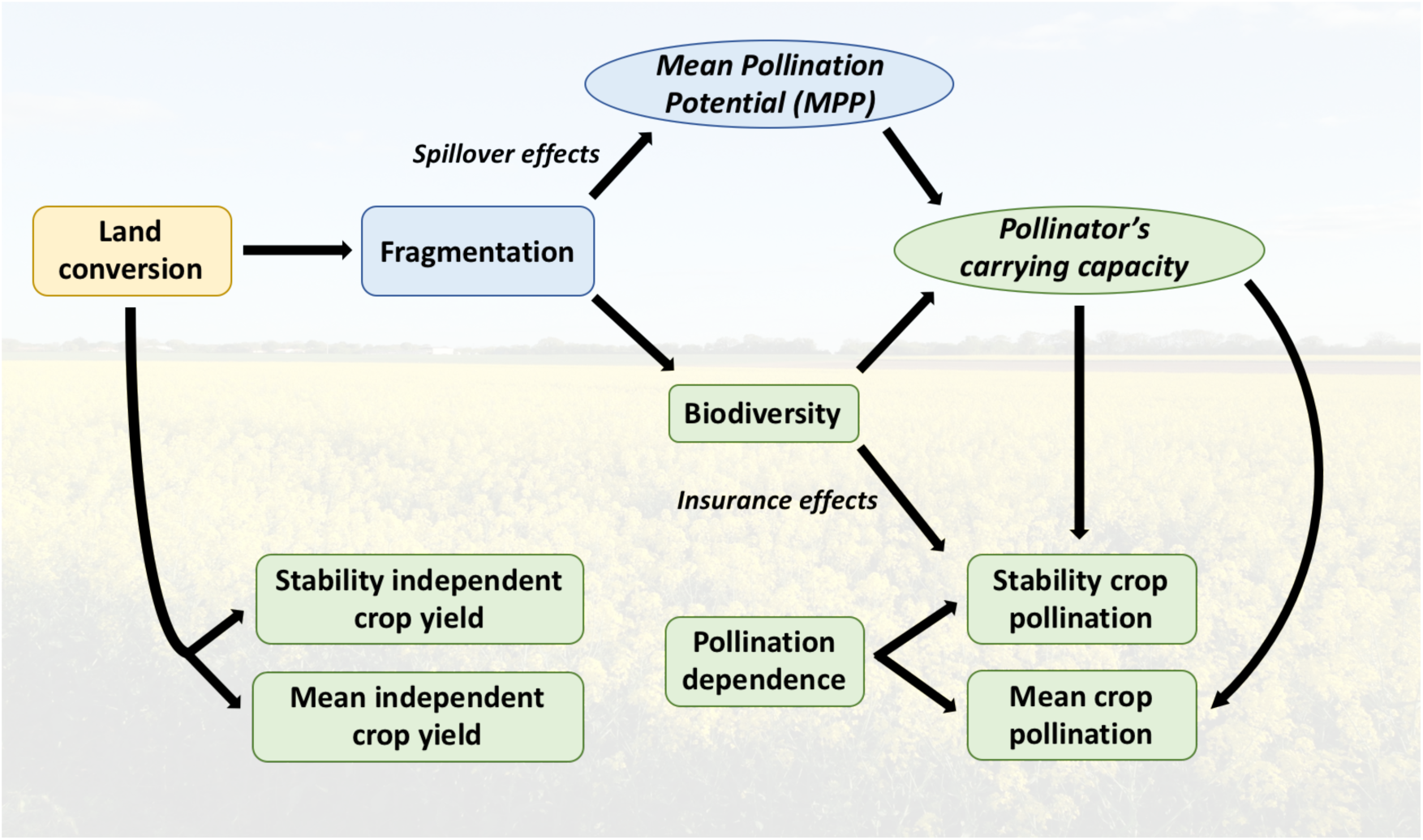
Conceptual diagram of our modelling framework. Green boxes represent non-spatial components of the model, whereas blue boxes are the additions that space brings to the model. Considering space allows: (i) to explore a continuous gradient of land conversion patterns, from completely random to highly aggregated, that encompasses a wide range of fragmentation scenarios, and (ii) to vary the strength of the spillover effect from seminatural habitat to crop land – i.e. the distance-decay of ecosystem service flows. Following a mean-field approximation, the *Mean Pollinator Potential* (MPP) of the agricultural landscape captures the full complexity of fragmentation effects of land conversion on ecosystem service supply that are not mediated by biodiversity (see results).

#### Crop dynamics

We used a model of biodiversity and crop biomass production in agricultural systems^20^, and extend it to a spatially-explicit landscape to investigate the response of ecosystem services to land conversion. The model represents intensively-managed agricultural landscapes, where crop land does not harbor significant levels of biodiversity. Spatial heterogeneity is defined by two types of patches: crop land and seminatural habitat. Crop land is used to grow annual crops with varying degrees of dependence on wild animal pollination, whereas seminatural habitat shelters biodiversity, including wild plants and pollinators. The model does not take managed honey bees into account as they do not depend on the availability of seminatural habitat, and they pollinate less efficiently compared to non-managed pollinators^31^. Crop land and seminatural habitat are linked by pollinators’ foraging movement. The three components of Montoya et al^20^ (pollinators, wild plants, and crop yield) are represented by the following equations (they have been transformed into their spatially-explicit, discrete-time versions):

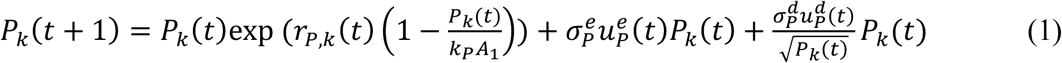

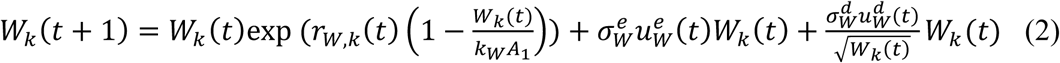

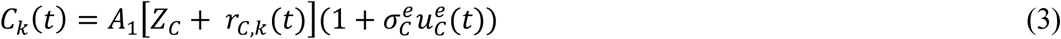

where *P*_*k*_ represents pollinators, *W*_*k*_ wild plants and *C*_*k*_ crop yield in cell *k*, with *P*_*k*_ (*t*) *= W*_*k*_ (*t*) *=* 0 if *k* is a crop land cell, and *C*_*k*_(*t*) *=* 0 if *k* is seminatural habitat. One unit of time *t* corresponds to one growing season, *P(t)* and *W(t)* can be interpreted as total biomass of pollinators and wild plants over growth season *t*, respectively, whereas *C(t)* is the total crop yield at the end of the growing season *t. L* is the set of crop land cells; *k*_*P*_ and *k*_*W*_ are the carrying capacities of pollinators and ‘wild’ plants, respectively, per unit area. *A*_*1*_ is the area of a single cell; *A* is total landscape area; *A*[1-*ω*_*sn*_] is the total crop land area, and *Aω*_*sn*_ is total seminatural area.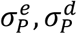 are the environmental and demographic stochasticity of pollinators, and 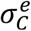 is the environmental stochasticity of crops. Equation (3), is the sum of pollination-dependent and independent parts of crop yield:

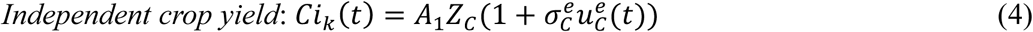

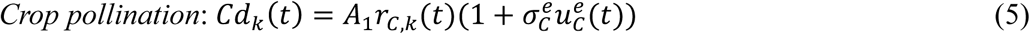

In the equations (1-3), *r*_*P,k*_ (*t*), *r*_*W,k*_ (*t*), *r*_*C,k*_ (*t*) are the pollinators’, ‘wild’ plants’ and crop’s per capita growth rates, and are defined as:

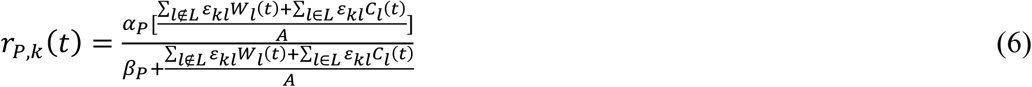

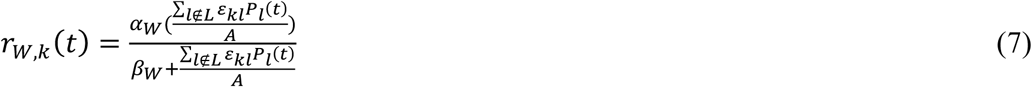

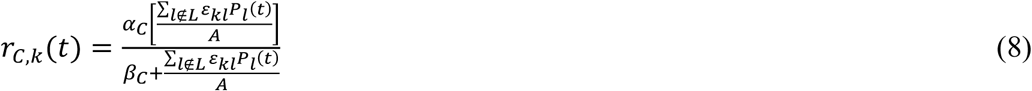

where *ε*_*kl*_ the distance-decay function representing the decrease of ecosystem service flow from seminatural habitat to crop land (see below). Pollinators are assumed to be generalist central-place foragers that feed on both wild plants and crops^32^, and pollinator’s growth rate thus depends on the availability of resources (wild plants and crops) in the neighborhood. The growth rate of wild plants does not depend on crops. Plant and pollinator uptake of resources follows a saturating, type II functional response, where *α*_P_ and *α*_W_ are the maximum growth rates; β_P_ and β_W_ are half-saturation constants. The pollination-dependent part of crop yield is also assumed to follow a type II functional response (Eq. 5 and 8), where *α*_C_ is the maximum crop yield derived from pollination, and β_C_ is the half-saturation constant of crops. The use of saturating functional responses is widely supported and consistent with real biological examples^33-35^. A complete description of the model parameters can be found in Table S1.

Ecosystem service provision at the landscape scale is estimated by summing up the individual contribution of each cell *k*. Thus, for total crop yield we obtained *C*(*t*) *=* ∑_*k∈L*_ *C*_*k*_(*t*). Crop yield per unit of agricultural area is calculated by dividing total crop yield by crop land area.

#### Distance-decay of ecosystem service flow

One main feature of space, as opposed to spatially-implicit or non-spatial systems, lies in the distance-decay of service flows from one habitat to another. For ecosystem service flow, we used a logistic distance-decay function adapted from Mitchell et al^19^. This function is consistent with both theoretical predictions^16,36-39^ and empirical observations^40-42^ of the effects of habitat edges and distance-to-habitat fragment gradients on ecosystem service provision. Other functions are possible and biologically meaningful (e.g. exponential decay), but they yield similar results (Mitchell et al^19^; Figure S1).

In our model, ecosystem service distance-decay mainly affects the flow of pollination to crop land. *ε*_*kl*_ is the distance-decay function of the effect pollinators on crops: the further a crop land cell is from seminatural habitat, the more difficult it is for pollinators to reach that cell, and thus the smaller the effect of *P*_*k*_ on crop biomass. To calculate *ε*_*kl*_, we adapt Mitchell et al^19^ distance-decay function as follows (see also Appendix S1):

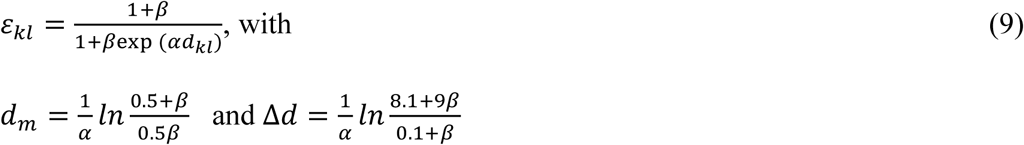

where *d*_*kl*_ is the distance between cells *k* and *l*, and parameters *α* and *β* are auxiliary parameters determined by

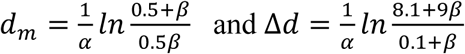

where *d*_*m*_ is the distance from seminatural cell edge at which the pollination flow equals one half of its initial value – 50% decay distance –, and Δ*d* is the distance over which the pollination flow decreases from 90% to 10% of its initial value – 90% to 10% decay distance. The total contribution of animal pollination to crop yield in a given cell *k* is made up from the summed contributions of individual seminatural habitat cells in the surrounding landscape. We explored a range of *d*_*m*_ values to investigate how variations in distance-from-fragment where service flow decreases influence ecosystem service supply. Variation in Δ*d* had little effect on model results (Figure S2; see also Mitchell et al^19^).

#### Stochasticity

To investigate the effects of land conversion pattern on yield stability, our model includes environmental and demographic stochasticity. Environmental stochasticity (e.g. variation in temperature, rainfall variability) is included through the terms *σ*^*e*^*u*^*e*^ *(t)*, where *(σ*^*e*^)^*2*^ is the environmental variance of either pollinators (*(σp*^*e*^)^*2*^)), ‘wild’ plants (*(σW*^*e*^)^*2*^) or crops (*(σC*^*e*^)^*2*^), and *u*^*e*^*(t)* are random functions with zero mean and standardized variance; we assume that perturbations have no temporal correlation. Demographic stochasticity (*σ*^*d*^*u*^*d*^ *(t)*) emerges from stochastic variation in individuals’ births and deaths. Crops are sown at high densities, and thus we assume demographic stochasticity is prevented in crops, and only affects pollinators and ‘wild’ plants. Demographic stochasticity is included in the form of the first-order normal approximation commonly used in stochastic population dynamics^43^, where *(σ*^*d*^)^*2*^ is the demographic variance of either pollinators (*(σ*_*P*_^*d*^)^*2*^) or ‘wild’ plants (*(σ*_*W*_^*d*^)^*2*^), and *u*^*d*^*(t)* are independent random functions with zero mean and standardized variance. For environmental stochasticity, we take the same perturbation for all cells and for all variables (because weather variations will be more or less the same over the entire landscape). For demographic stochasticity, we take independent perturbations between cells and variables.

#### Biodiversity and fragmentation

Despite recent debate has ensued on the relative importance of habitat loss *versus* fragmentation on species diversity^15,44-46^, empirical evidence shows that larger and more connected fragments of natural habitat in general host more biodiversity than smaller and more isolated fragments^14^. In agricultural landscapes, this means that different land conversion patterns (e.g. random, aggregated) will result in different biodiversity levels which will in turn influence ecosystem service supply in many ways. Hanski et al^47^ proposed a way to capture the relationship between biodiversity and habitat fragmentation, namely the *Species-Fragmented Area Relationship* (SFAR), which extends the conventional species-area relationship (SAR) to landscapes where fragmentation pervades. The SFAR has the following form:

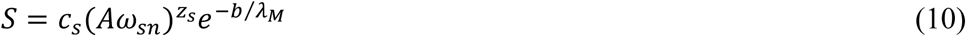

where *S* is pollinator diversity – species richness –, and *Aω*_*sn*_ is the total area of seminatural habitat; *b* is a parameter modulating the effect of the metapopulation capacity and reflects the ability of species to live in fragmented landscapes (e.g. low *b* characterized species evolved or well adapted to live in fragmented landscapes). The degree of fragmentation is captured by *λ*_*M*_, which represents the metapopulation capacity of the fragmented landscape. The metapopulation capacity *λ*_*M*_ is obtained from the leading eigenvalue of a *n* x *n* matrix with elements *m*_*ii*_ = 0 and 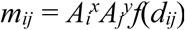, where *A*_*i*_ and *A*_*j*_ are the areas of fragments *i* and *j, x* and *y* are scaling factors (we use *x* = 2, *y* = 1 as in Hanski et al^47^), *d*_*ij*_ is the Euclidean distance between the centroids of fragments *i* and *j*, and *f*(*d*_*ij*_) is the dispersal kernel. Following Hanski et al (2013), we assume the exponential dispersal kernel with a cutoff at 0.01, *f*(*d*_*ij*_) = max{exp(-*δd*_*ij*_), 0.01}, where 1/*δ* gives the average dispersal distance, and estimated *λ*_*M*_ from information on fragment size and distance among fragments (all referred to seminatural habitat). We used the accepted value of *z*_*s*_ = 0.25 for a wide range of plants and animals^48^, and allowed dispersal distance and *b* to vary.

Changes in landscape structure can affect biodiversity and the ecosystem functions that underlie ecosystem service provision. To consider the effects of fragmentation on biodiversity and crop pollination, we made crop pollination dependent on pollinator diversity. This was done by creating a dependence of pollinator’s carrying capacity (*k*_*P*_) on biodiversity following a power law: 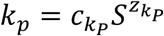, where *S* is the number of pollinator species estimated by the SFAR, and 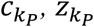 are the parameters of the power law. We use the values of 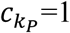 and 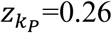 based on recent literature^49-51^, but also considered two extreme values of *h* (0.0, 0.5) to more clearly explore the effect of pollinator diversity. Finally, we considered the ability of pollinator diversity to provide an insurance against environmental fluctuations, i.e. insurance effect of biodiversity. To do this, we made 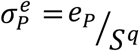 and compared a scenario where environmental stochasticity depends on biodiversity (*q* = 1/2)^52^ with another scenario where biodiversity does not affect environmental stochasticity (*q =* 0). Demographic stochasticity acts at the individual level, and in the same manner for conspecifics and heterospecifics; thus, there is no insurance effect for demographic stochasticity.

### b. Land conversion pattern generation

The landscape consisted of two-dimensional lattice (25 × 25 cells) where individual cells can have either of two states: crop land or seminatural habitat. We generated land conversion patterns by iteratively creating crop land cells in a landscape that consisted initially only of seminatural land. In a single step of the algorithm only one semi-natural habitat cell is selected and converted. At each iteration, we determined for each seminatural land cell the number of neighboring crop land cells, a number we denote by *m* (*m* is equal to 0, 1, 2, 3 or 4). We then chose randomly one of the seminatural land cells, with a probability that depended on the number of neighboring crop land cells. More precisely, the probabilities were proportional to *p* = 0.1^*w*^ if *m* = 0 and *p* = *m*^*w*^ if *m* ≥ 1. These values are actually relative probabilities; that is, they have to be normalized to get the probability of selecting a given cell. Hence, for *w* = 0 all seminatural land cells had the same relative probability to be chosen, leading to a fully random, unclustered pattern. For *w* > 0, seminatural land cells with more neighboring crop land cells had a higher relative probability to be converted, leading to a clustered or aggregated pattern. Larger values of *w* resulted in more aggregated patterns. Therefore, variation in the value of *w* allowed us to produce a continuous gradient of land conversion patterns, and therefore fragmentation, based on the aggregation degree (Figure S3). For each land conversion pattern, we characterised fragmentation of the remaining seminatural habitat by quantifying mean fragment size, number of fragments, mean fragment perimeter, and perimeter: area ratio.

## RESULTS

### Mean-field approximation

Because the spatially explicit model demands much computational time, we analysed how the spatially-explicit model is linked to the spatially-implicit one. To do so, we developed a mean-field approximation of the spatial agroecosystem model (Eqs. 1-3), which replaces the detailed spatial structure of the landscape by a much simpler, spatially averaged one (see Appendices S2 and S3 for further details on the solution of the full model and the derivation of the mean-field approximation). To do this, consider the sums ∑_*l*∉*L*_ *ε*_*kl*_ (Eq. 8): there are *ω*_*sn*_*n*^*2*^ terms (possible values of *l*; *n*^*2*^ is the number of cells in the agricultural landscape), and (1-*ω*_*sn*_)*n*^*2*^ such sums (possible values of *k*). It turns out that the main effects of the spatial structure can be accounted for by a new parameter,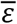, defined as

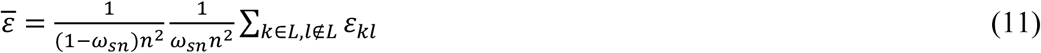

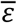 is the average value of *ε*_*kl*_ when taking a random cell *k* ∈ *L* and a random cell 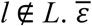 has two complementary interpretations. Firstly, it is a measure of the amount of seminatural habitat supplying pollinators to crop land: if we multiply 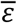 by the area of seminatural habitat 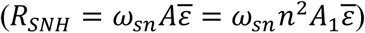, then *R*_*SNH*_ is the area from which a crop land cell can be pollinated averaged over all crop land cells. Secondly, 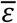 quantifies the amount of crop land that is reachable by pollinators from seminatural habitat: if we multiply 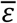 by the crop land area 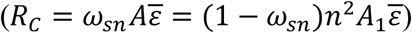, *R*_*C*_ is the crop land area that a pollinator can reach averaged over all seminatural cells. Taken together, these two interpretations can be summarized by the term *Mean Pollination Potential* (MPP; 0 ≤ MPP ≤ 1). Under the mean-field approximation the mean crop yield is (see Appendix S3):

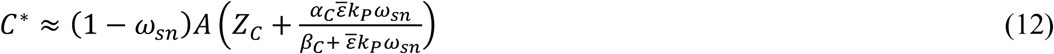

The variance of crop yield is:

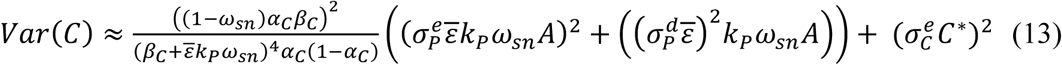

with *k*_*p*_ *= aS*^*h*^ (see methods). We found that the mean-field approximation is a very accurate description of the dynamics of various ecosystem services in agricultural landscapes, both for mean and stability values (Figure 2).

**Figure 2.**
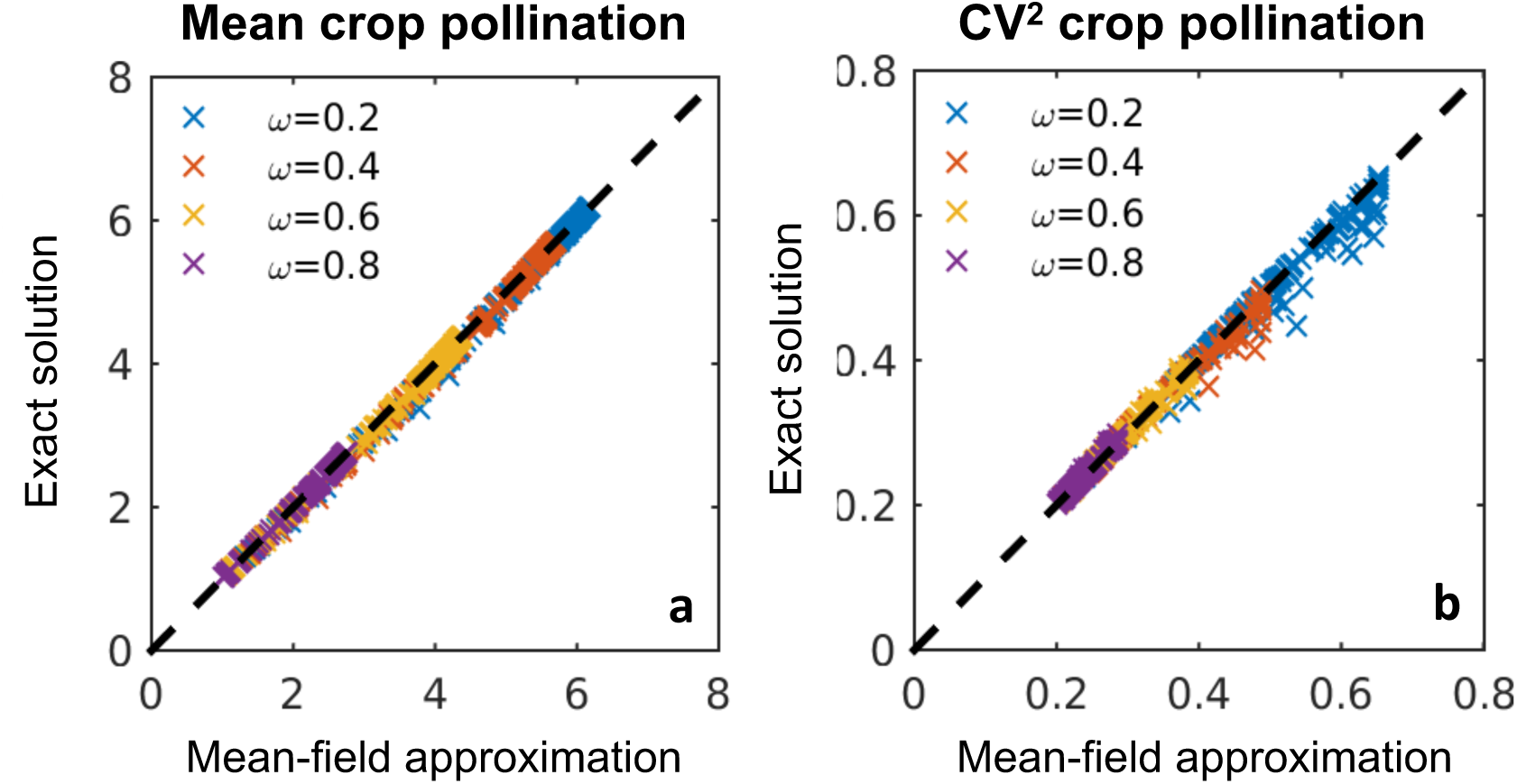
Mean-field approximation vs Exact solution. (**a**) Mean crop pollination. (**b**) Variability of crop pollination (measured as Coefficient of Variation – CV –, the inverse of stability). Exact solution equations can be found in Appendix 2 (Eqs. 8 and 17 for crop pollination mean and variability, respectively). Mean-field results are derived from Eqs. 12 and 13 in the main text, for mean and variability of crop pollination, respectively. *ω*_*Sn*_ is the proportion of seminatural habitat (drawn randomly in [0,5], Figure S3). *d*_*m*_ = drawn randomly in [1,25], expressed in linear dimension of a landscape cell, 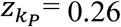. Parameter values: α_P_ = α_W_ = 0.9, β_P_ = β_W_ = 0.6, *A* = 10, *Z*_*C*_ = 1000, α_C_ = 1000, *k*_*W*_ = 5000, *k*_*P*_ = 0.1, e_P_ = 0.8, σ^d^_P_ = 0.1, σ^e^_C_ = 0.03, α_C_ = 1000, Pollination dependence = 50%.

The mean-field approximation shows that the fragmentation effects of land conversion on crop pollination services are determined by MPP. To consider the spatial structure of land conversion, the term *β*_*C*_/*k*_*P*_ of the non-spatial model^20^ has to be replaced by

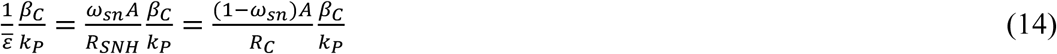

*β*_*C*_/*k*_*P*_ is the ratio of crop half-saturation constant relative to pollinators’ carrying capacity, and is an *effective* parameter combination that strongly influences crop dynamics, as it quantifies the pollinator requirement of crops relative to the availability of pollinators, i.e. crop relative requirement for pollinators. When *β*_*C*_/*k*_*P*_ is small, crop yield saturates at lower pollinator biomass than their carrying capacity; when *β*_*C*_/*k*_*P*_ is large, crop yield saturates at pollinator biomasses much higher than their carrying capacities. *β*_*C*_/*k*_*P*_ influences both the mean and stability of crop pollination. On one hand, greater values of *β*_*C*_/*k*_*P*_ increase the effect of pollinator biomass on crop pollination, reducing mean yield and shifting maximum yield to larger amounts of seminatural habitat. On the other hand, *β*_*C*_/*k*_*P*_ controls how fast the saturation of crop pollination to pollinator biomass sets in and, consequently, how fast the response of crops to pollinator stochasticity drops down; thus, the smaller *β*_*C*_/*k*_*P*_ the faster the saturation sets in, and so the faster crop yield variability drops when increasing seminatural habitat (Figure S4A). Without distance-decay (or when MPP ≈ 1), the spatial model collapses into the non-spatial model (Figure 3A-C, dark blue lines; Figure S4B). Fragmentation effects on ecosystem services become stronger when MPP<1, which increases *β*_*C*_/*k*_*P*_.

**Figure 3.**
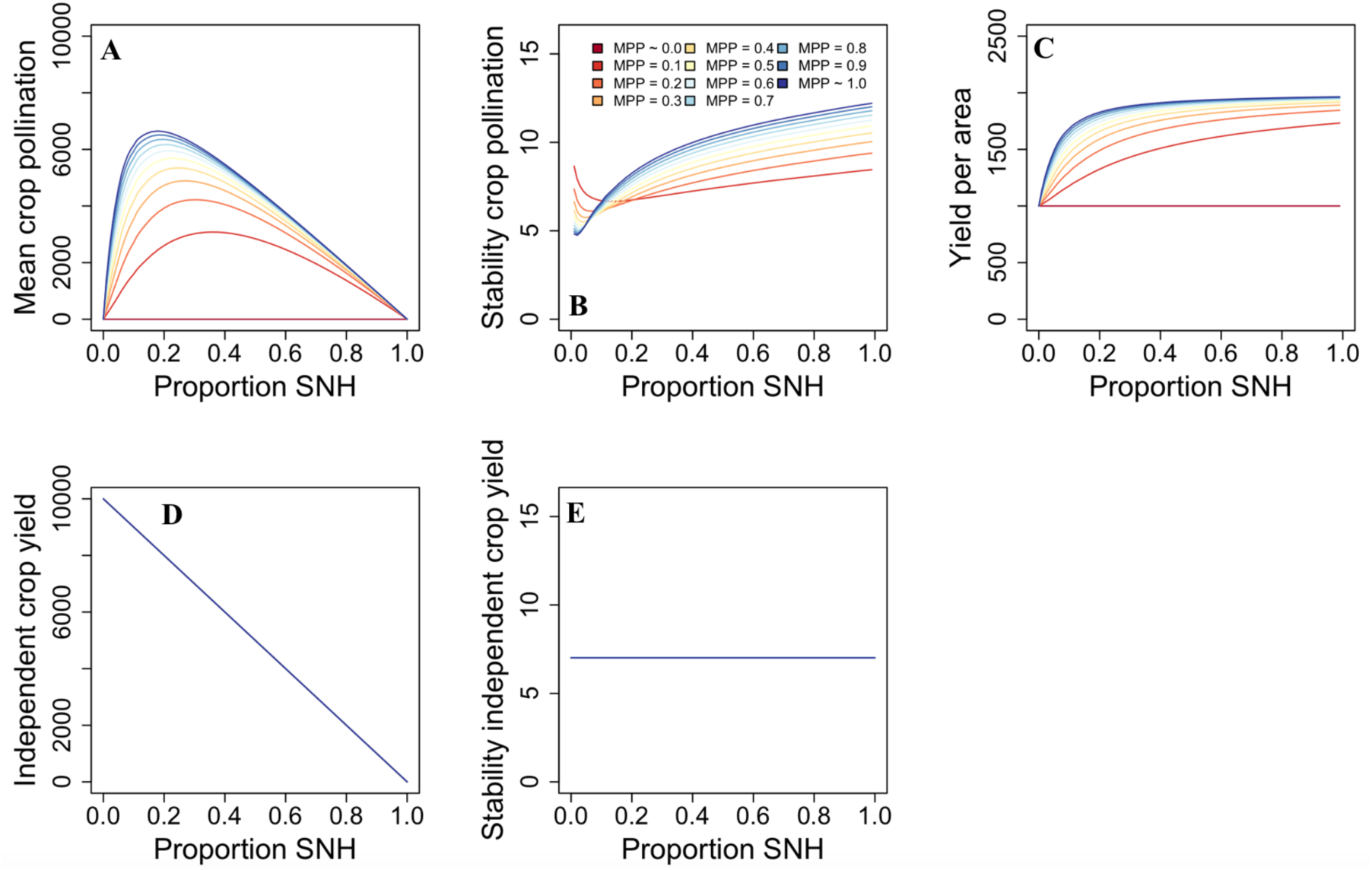
Effects of landscape composition and MPP on ecosystem services. Ecosystem services are represented as a function of the proportion of seminatural habitat, for different MPP. MPP includes the effects of fragmentation – more specifically, the aggregation pattern of land conversion – and the distance-decay of ecosystem service flow. Parameter values: α_P_ = α_W_ = 0.9, β_P_ = β_W_ = 0.6, *A* = 10, *Z*_*C*_ = 1000, α_C_ = 1000, *k*_*W*_ = 5000, e_P_ = 0.8, σ^d^_P_ = 0.1, σ^e^_C_ = 0.03, α_C_ = 1000, Pollination dependence = 50%, 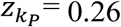.

### Spatial constraints/ fragmentation effects on MPP

MPP depends on two factors: fragmentation – more specifically, the aggregation pattern of land conversion – and the distance-decay of ecosystem service flow. High aggregation (low fragmentation) and fast distance-decay result in lower MPP (Figures S5 and S6), which in turn reduce crop pollination services. These two factors interact: only when the flow of pollinators to crop land is limited (fast distance-decay) aggregation patterns are relevant for crop production (Figure S5A-D). In this case, higher aggregation, through its effects on MPP, not only reduces mean crop pollination and shifts maximum yield to higher fractions of seminatural habitat, but also decreases yield stability along the gradient of seminatural habitat (Figure 3A, B). When no restrictions exist in the flow of pollinators to crop land, MPP is maximum (MPP ≈ 1; Figure 3) and fragmentation does not affect pollination services (Figure S5E-F).

### MPP effects on ecosystem services

We did not find any clear, consistent effect of specific fragmentation metrics on ecosystem services (Figure S7). However, the full complexity of the purely spatial fragmentation effects (i.e. those not mediated by biodiversity) on ecosystem service supply, irrespective of the specific pattern of land conversion, were captured by MPP (Figure S8). When MPP =1, fragmentation effects are negligible and crop dynamics are identical to those of the non-spatial model (Figure 3A-C, dark blue lines). In this case, no additional mechanisms need to be invoked: crop yield dynamics are driven by the crop’s relative requirement for pollinators (*β*_*C*_/*k*_*P*_, Figure S4) and the degree to which crops depend on animal pollination. The effects of fragmentation kick off when MPP < 1. Lower MPP – i.e. more aggregated patterns of land conversion (Figure S5) – reduces the carrying capacity of pollinators (Eq. 14), which decreases the provision of pollinator-dependent ecosystem services (Figure 3A-C). The same is true for crop pollination stability, except at small fractions of seminatural habitat and/or small values of MPP. A higher biodiversity effect 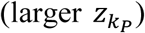 increases both mean crop pollination and its stability, as well as yield per area (Figure S9). MPP has no effect on independent crop yield as it does not depend on animal pollination and, therefore, on seminatural habitat (Figure 3D-E).

The effect of MPP on crop pollination services increases with the degree to which crops depend on animal pollination. Higher pollination dependence of crops shifts maximum yields to higher fractions of seminatural habitat at landscape and local scales, and the stability of crop pollination increases faster (Figure S10).

### Biodiversity effects on crop pollination

Biodiversity decreases with land conversion, but higher aggregation of seminatural fragments alleviates that loss to some extent (Figure S11). The effects of fragmentation on biodiversity are stronger at low-intermediate fractions of seminatural habitat, and are directly influenced by the dispersal distance of organisms and by their ability to live in fragmented landscapes (Figure S12). Biodiversity stabilizes crop pollination by increasing the pollinators’ carrying capacity (which affects the variance of crop production, Eq. 13; Fig. 4B), and by reducing the response of crop pollination to environmental fluctuations (Fig. 4C). The former effects are stronger when biodiversity is higher, whereas the latter effects reduce variability of crop pollination especially at increasing biodiversity levels. A higher biodiversity effect 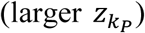 increases both mean crop pollination and its stability, as well as yield per area (Figure S13).

**Figure 4.**
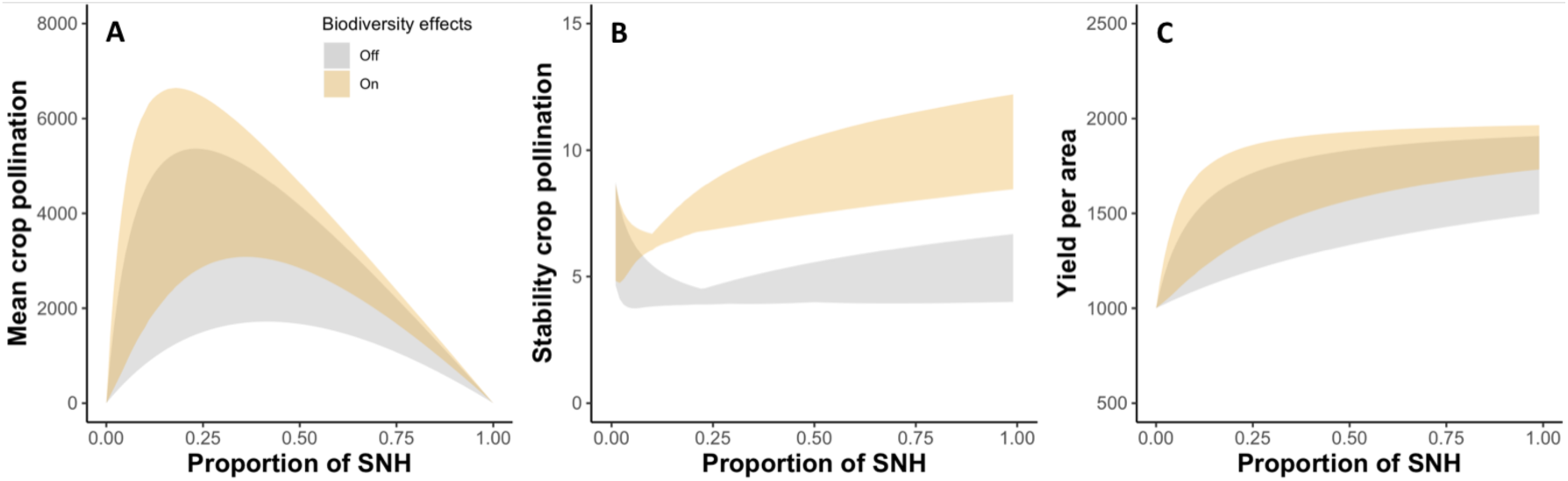
Effects of biodiversity on crop pollination. Plots show the response of crop pollination services – mean and stability of crop pollination (panels **A** and **B**), and yield per area (panel **C**) – as a function of the proportion of seminatural habitat (SNH). All MPP values are contained within the shadows, whose limits are determined by the minimum and maximum values across the range of MPP. Biodiversity can affect crop pollination in a two-way manner. On one hand, biodiversity influences mean crop pollination and yield per area by increasing the carrying capacity of pollinators 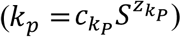. On the other hand, biodiversity impacts the stability of crop production both indirectly – increasing the carrying capacity of pollinators – and directly – reducing the response of crop production to environmental fluctuations 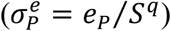. For each ecosystem service, the plots compare two scenarios: (i) a scenario where biodiversity has no effect on crop pollination (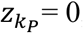, *q* = 0), represented by the grey shadows, *versus* (ii) a scenario where biodiversity has an effect on crop pollination (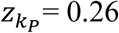, *q* = ½; Tilman 1999, Liang et al 2016, O’Connor et al 2017, Cardinale et al 2011), represented by the light orange shadows. Parameter values: α_P_ = α_W_ = 0.9, β_P_ = β_W_ = 0.6, *A* = 10, *Z*_*C*_ = 1000, α_C_ = 1000, *k*_*W*_ = 5000, e_P_ = 0.8, σ^d^_P_ = 0.1, σ^e^_C_ = 0.03, α_C_ = 1000, Pollination dependence = 50%.

### Net effects of fragmentation on ecosystem services

The fragmentation pattern of seminatural habitat has a dual effect on crop pollination services. On one hand, aggregation of seminatural fragments decreases pollination by lowering MPP (Figures S5 and S14), which in turn reduces the carrying capacity of pollinators (Eq. 14). On the other hand, aggregation increases biodiversity (especially at low-intermediate fractions of seminatural habitat; Figure S11), which in turn increases pollinators’ biomass (through its positive effects on pollinators’ carrying capacity) and the service of pollination (Figure S14). The net effect of fragmentation on ecosystem service supply depends on the distance-decay of ecosystem service flow (*d*_*m*_) and the proportion of seminatural habitat remaining. When the decay distance *d*_*m*_ is low (Figure 5, first row), fragmentation effects tend to be positive for mean crop pollination and yield per area because the fraction of crop land within reach from non-crop land areas is higher (this fraction is lower at very low fractions of seminatural land). Yet, crop pollination stability decreases due to the lower biodiversity levels in fragmented landscapes, except at high fractions of seminatural habitat where the impact of fragmentation is minimum. Conversely, when the decay distance *d*_*m*_ is high, seminatural fragments are perceived as more connected and ecosystem service supply is not limited by space. In this case, fragmentation becomes irrelevant, or even negative, due to the lower biodiversity levels in fragmented landscapes (Figure 5).

**Figure 5.**
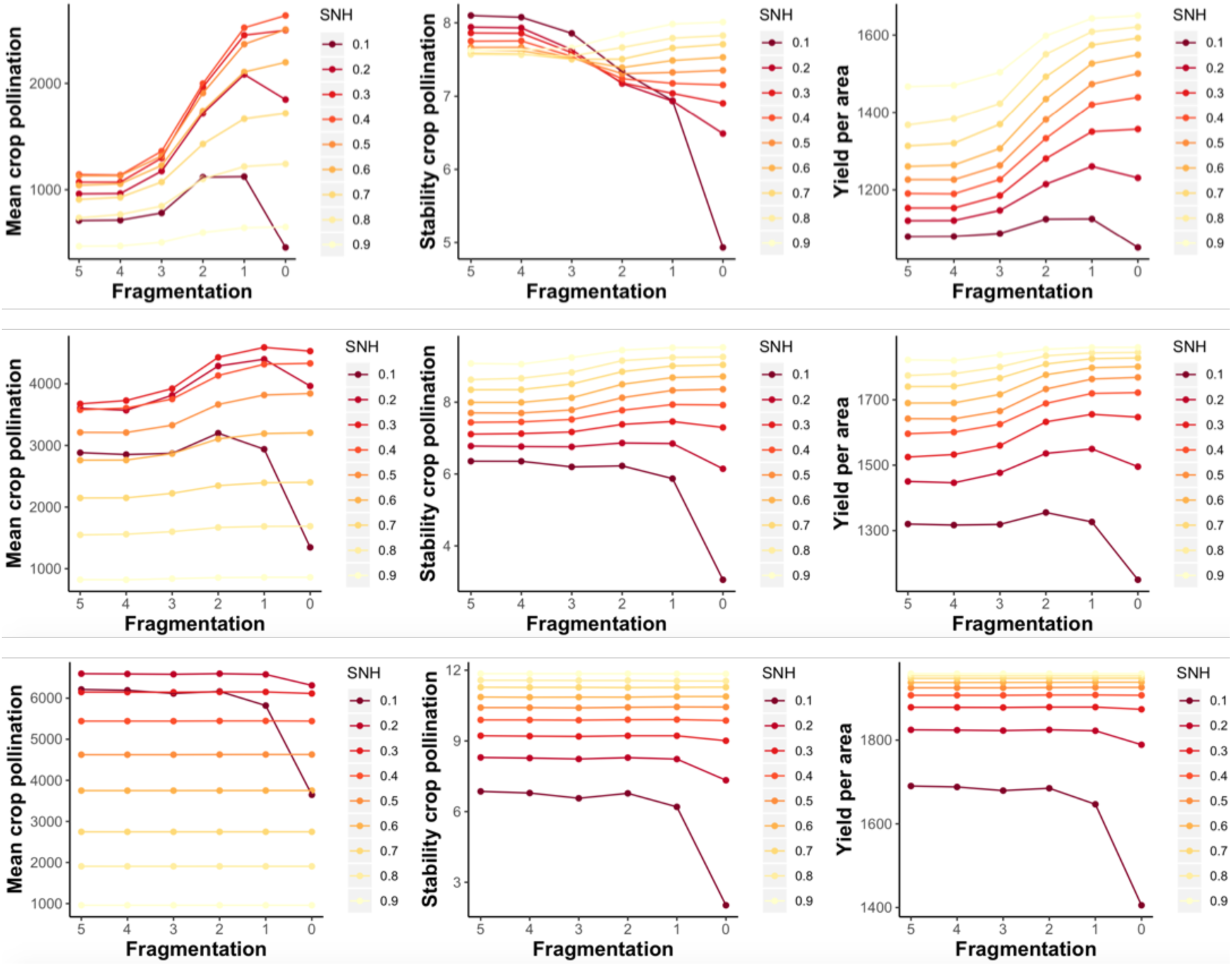
Net effects of aggregation on crop pollination services. Columns represent, from left to right, mean and stability of crop pollination, and yield per area. Ecosystem services are plotted as a function of fragmentation for different proportion of seminatural habitat or SNH (as opposed to figures 3-4). Fragmentation increases in the x-axis from left to right (we set *w* = *m* for simplicity; higher *w, m* means more aggregation). Darker lines correspond to lower fractions of seminatural habitat, which are more typical of intensive farming systems. Rows represent increasing values of the decay distance *d*_*m*_ (0.5, 1, 5). Parameter values: α_P_ = α_W_ = 0.9, β_P_ = β_W_ = 0.6, *A* = 10, *Z*_*C*_ = 1000, α_C_ = 1000, *k*_*W*_ = 5000, e_P_ = 0.8, σ^d^_P_ = 0.1, σ^e^_C_ = 0.03, α_C_ = 1000, Pollination dependence = 50%, 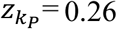.

## DISCUSSION

Our analysis reveals a variety of effects of land conversion on biodiversity and crop production in intensive crop pollination systems. Using a mean-field approximation of various ecosystem services in spatially-explicit agricultural landscapes, our model suggests that (1) fragmentation impacts food production through spatial and biodiversity-mediated effects; (2) the full complexity of the fragmentation-induced spatial effects on ecosystem service supply, irrespective of the specific pattern of land conversion, is captured by one factor – the mean pollination potential of the remaining seminatural land (MPP) – which determines the mean and stability of pollination services; (3) biodiversity can have a stabilizing effect on crop pollination in fragmented agricultural landscapes; and (4) the net effects of fragmentation on food production depend on the strength of the spillover of pollinators to crop land and the degree to which crops depend on animal pollination.

The loss of seminatural land has contrasting effects on the ecosystem services considered: biodiversity decreases, independent crop production increases, while crop pollination is maximized at intermediate fractions of seminatural habitat. But fragmentation can modify these relationships in two ways. On one hand, land conversion can produce multiple patterns of aggregation of the remaining fragments of seminatural habitat. These patterns combined with the strength of the spillover of pollinators to crop land determine the mean pollination potential of seminatural land (MPP), which is the main responsible of food production in pollination-dependent agriculture. The second type of effects are mediated by biodiversity, as the level of aggregation of the remaining fragments of seminatural habitat affects the pollinator richness. Such purely spatial and biodiversity-mediated effects modify the carrying capacities of pollinators, which ultimately determine crop pollination services. The mean-field approximation shows that the effects of space on crop production can be interpreted in the same terms as varying the pollinator’s carrying capacity in the non-spatial model^20^.

Our results suggest that understanding the factors that affect MPP is a fundamental step towards food security. If no restrictions exist in the flow of pollinators to crop land, MPP is maximum and the spatial structure of land conversion does not affect crop yield dynamics. In this situation, seminatural fragments are perceived as more connected and the provision and stability of crop pollination is not conditioned by space, i.e. spatial and non-spatial models converge. However, agricultural landscapes are fragmented to some extent and the foraging ranges of most organisms are local (200 m for small bee species, 25–110 m for bumble bees, >200 m for certain bee species^53-56^), which produces higher aggregation and weaker spillover effects, thus reducing MPP. Such reductions in MPP affect crop yields by (i) decreasing mean crop pollination and total yield per area, and (ii) decreasing yield stability along the gradient of seminatural habitat. The estimation of MPP in real farming systems would require data on the aggregation level of seminatural habitat fragments within the agricultural landscape, and on the spillover of pollinators to adjacent crops. The former can be obtained with GIS processing of aerial pictures or satellite images of agricultural landscapes. For the latter, information on foraging distances of pollinator species combined with experimental studies could be used to reveal species’ foraging patterns and how the flow of pollinators to adjacent crop land decays with distance (e.g.^16,36,37-39^). This information will be useful to design agricultural landscapes for high MPP.

Producing food requires land, and increasing the land devoted to farming reduces the land devoted to biodiversity conservation. Our results agree with recent empirical studies showing that higher pollinator diversity increases food production^6^, and further suggest that it can lead to lower variability in agricultural productivity. The response of biodiversity to land conversion depends on the amount and the spatial structure of seminatural habitat loss. For example, although the effects of fragmentation on biodiversity are stronger at low-intermediate fractions of seminatural habitat – typical of intensive farming systems –, aggregation increases the biodiversity levels within seminatural habitat fragments. The stabilizing effect of biodiversity and its role in food security is increasingly supported, even at crop levels^12^. Our results add to this view and point to biodiversity conservation as one key policy to achieve food security.

Our findings are consistent with previous studies that found non-linear effects of fragmentation on ecosystem service provision (e.g.^18,19^), and provide a theoretical basis of the effects of fragmentation on the stability patterns of crop pollination. Fragmentation has a dual effect on crop production services. On one hand, aggregation decreases crop pollination by reducing MPP. On the other hand, aggregation increases crop pollination by maintaining higher biodiversity, especially at low-intermediate fractions of seminatural habitat. The net effects of aggregation on crop pollination depend on the strength of spillover effects. These results have management implications (e.g. land sharing–sparing debate^57,58^), as the goals of different landscape managers can be conditioned by the way that natural land is converted into crops. For example, maintaining a large number of seminatural fragments may be a better strategy at multiple spatial scales than maintaining a few large fragments when pollinator flow to crop land is low. Yet, this strategy may increase the temporal variability of crop pollination at low-intermediate proportions of seminatural habitat, reflecting a trade-off between ecosystem service mean and stability. Conversely, larger fragments of seminatural habitat have higher pollinator diversity when the fraction of seminatural habitat is low or intermediate, and higher biodiversity can stabilize crop pollination. These results agree with recent claims that the land sharing–sparing dichotomy lends itself to overly simplistic policy prescriptions^59^, and suggest that management decisions for food security should consider factors such as the distance-decay of pollinator flow, the amount and spatial aggregation of seminatural habitat and the degree to which crops depend on animal pollination.

Our model has several limitations. For example, our model focuses on intensive farming systems, where crop land does not host important biodiversity levels; other types of agriculture – e.g. organic farming, wildlife-friendly practices – allow moderate levels biodiversity to thrive within crop land, and can modify the results reported here^60^. Second, the observation that biodiversity loss has either none (stability) or positive (mean) effects on independent crop yield may change if organisms responsible for other services, i.e. pest control, are included. Besides, although we do not find any effect of seminatural habitat on the stability of independent crop yield, this may change if environmental stochasticity of crops increases with decreasing amounts of seminatural habitat, as suggested by studies linking seminatural habitat to climate regulation, natural hazard regulation and water flow regulation services^61^. Finally, our model focuses on wild central-place pollinators (i.e. all types of wild bees, including bumble bees and solitary bees), whose presence and abundance directly depend on the amount of seminatural habitat^32^, which provides shelter and habitat for these insects. Honey bee colonies are used to substitute wild pollinator communities, yet the pollination services of wild pollinators cannot be compensated by managed bees because (1) pollinator-dependent crop land grows more rapidly than the stock of, e.g., honey bee colonies^62^, (2) wild insects usually pollinate crops more efficiently than honey bees^31^, and (3) honey bees may depress wild pollinator densities^63^. Despite other groups of pollinators exist, wild central-place foragers remain a very important group of crop pollinators in agriculturally dominated landscapes^64,65^.

Ensuring stable food supplies is one of the 2017 UN Sustainable Development Goals, and is a challenge that may require multiple solutions. Policies to increase yields, changing diets, irrigation, crop diversity, tolerance of crops to drought, among others, have been proposed as stability-enhancing solutions^12,27,66-68^. Our study sheds new light in this debate by showing that high and stable yields in crop pollination systems depend on biodiversity and the spatial structure of the agricultural landscape, i.e. fragmentation. Fragmentation can produce spatial and biodiversity-mediated effects with the potential to modify the mean and stability of pollination-dependent crop production, which has strong consequences for food production and food security. These results are highly relevant given the worldwide trends in agriculture, which shifts towards more pollinator-dependent crops^29,30^.

## Acknowledgements

DM was funded by the EU and INRA in the framework of the Marie-Curie FP7 COFUND People Program, through the award of an AgreenSkills/AgreenSkills+ fellowship, and by and the FRAGCLIM Consolidator Grant, funded by the European Research Council under the European Union’s Horizon 2020 research and innovation programme (grant agreement number 726176). This work was supported by the TULIP Laboratory of Excellence (ANR-10-LABX-41) and by the BIOSTASES Advanced Grant funded by the European Research Council under the European Union’s Horizon 2020 research and innovation program (grant agreement no. 666971).

## Author contributions

D.M., B.H. and M.L. conceived the original idea and designed the research. D.M. and B.H. designed the model, with help from M.L. and C.M. D.M. and B.H. performed the analysis. D.M. wrote the first draft of the manuscript, all authors contributed to revisions.

## Additional information

Supplementary Methods (3)

Supplementary Tables (1)

Supplementary Figures (15)

