## Supplementary Material for "Habitat fragmentation and food security in crop pollination systems"

### Supplementary Methods 1. Distance-decay function

Mitchell et al (2015) propose the following function to describe how the provision of an ecosystem service decreases with spatial distance,

$$\tilde{\varepsilon}(d) = 1 - \frac{1}{1 + \exp\big(-\frac{\ln(81)}{\Delta d}(d - d_m)\big)}$$

This function, which is monotonically decreasing, takes the following values:

$$\begin{aligned}\tilde{\varepsilon}(d_m - \tfrac{\Delta d}{2}) &= 0.9\\ \tilde{\varepsilon}(d_m) &= 0.5\\ \tilde{\varepsilon}(d_m + \tfrac{\Delta d}{2}) &= 0.1\end{aligned}$$

However, it does not equal 1 at $d = 0$, as one would expect naively.

The question is then, can we construct a modified function $\varepsilon(d)$ which has the same "logistic" shape as $\tilde{\varepsilon}(d)$ and satisfies $\varepsilon(0) = 1$. We start by rewriting $\tilde{\varepsilon}(d)$,

$$\begin{aligned}\tilde{\varepsilon}(d) &= 1 - \frac{1}{1 + \exp\big(-\frac{\ln(81)}{\Delta d}(d - d_m)\big)}\\ &= \frac{\exp\big(-\frac{\ln(81)}{\Delta d}(d - d_m)\big)}{1 + \exp\big(-\frac{\ln(81)}{\Delta d}(d - d_m)\big)}\\ &= \frac{1}{1 + \exp\big(\frac{\ln(81)}{\Delta d}(d - d_m)\big)}\\ &= \frac{1}{1 + \exp\big(-\frac{\ln(81)}{\Delta d}d_m\big)\exp\big(\frac{\ln(81)}{\Delta d}d\big)}\end{aligned}$$

showing that it has the functional shape

$$\tilde{\varepsilon}(d) = \frac{1}{1 + \beta\exp(\alpha d)}$$

for parameters $\alpha$ and $\beta$. If we want to keep the same shape for $\tilde{\varepsilon}(d)$, and additionally impose that $\varepsilon(0) = 1$, it suffices to set

$$\varepsilon(d) = \frac{1 + \beta}{1 + \beta\exp(\alpha d)}$$

We have

$$\begin{aligned}\varepsilon(d_1) &= 0.9 \qquad \text{for} \qquad d_1 = \frac{1}{\alpha}\ln\frac{0.1 + \beta}{0.9\beta}\\ \varepsilon(d_2) &= 0.5 \qquad \text{for} \qquad d_2 = \frac{1}{\alpha}\ln\frac{0.5 + \beta}{0.5\beta}\\ \varepsilon(d_3) &= 0.1 \qquad \text{for} \qquad d_3 = \frac{1}{\alpha}\ln\frac{0.9 + \beta}{0.1\beta}\end{aligned}$$

so that the relationship between $(\alpha, \beta)$ and the parameters $(d_m, \Delta d)$ of Mitchell et al (2015) is given by

$$d_m = \frac{1}{\alpha}\ln\frac{0.5 + \beta}{0.5\beta} \qquad \text{and} \qquad \Delta d = \frac{1}{\alpha}\ln\frac{8.1 + 9\beta}{0.1 + \beta}$$

Note however that the additional distance to decay from 0.9 to 0.5 and the additional distance to decay from 0.5 to 0.1 are no longer equal, as was the case for the function $\tilde{\varepsilon}(d)$ of Mitchell et al (2015).

It is interesting to note that the exponential function is a special case of $\varepsilon(d)$. To see that, take a large value for $\beta$, so that $\varepsilon(d) = \exp(-\alpha d)$, $d_m = \ln(2)/\alpha$ and $\Delta d = \ln(9)/\alpha$.

### Supplementary Methods 2. Equilibrium and variability of pollinator biomass, wild plant biomass, and crop yield

Here we derive for the discrete-time dynamics the expressions for the equilibrium and variability of pollinator biomass $P$, wild plant biomass $W$ and crop yield $C$.

#### Model equations – deterministic part

The deterministic part of the dynamics for pollinator biomass $P$ and wild plant biomass $W$ obey the following ordinary differential equations (ODEs),

$$P(t+1) = P(t)\exp\Big(r_P\big(1 - \frac{P(t)}{k_P\omega_{\rm sn}A}\big)\Big) \tag{1}$$

$$W(t+1) = W(t)\exp\Big(r_W\big(1 - \frac{W(t)}{k_W\omega_{\rm sn}A}\big)\Big), \tag{2}$$

with

$$r_P = c_P\frac{\alpha_P\big(\phi_W W(t) + \phi_C C(t)\big)}{\beta_P + \phi_W W(t) + \phi_C C(t)} \tag{3}$$

$$r_W = c_W\frac{\alpha_W P(t)/A}{\beta_W + P(t)/A}. \tag{4}$$

Crop yield $C$ is modeled as an instantaneous function (i.e., no dynamics) of pollinator biomass $P$,

$$C(t) = (1-\omega_{\rm sn})A\Big(Z_C + \frac{\alpha_C P(t)/A}{\beta_C + P(t)/A}\Big). \tag{5}$$

#### Equilibrium solution

The equilibrium values of pollinator biomass $P$ and wild plant biomass $W$ are readily obtained by setting the right-hand side of equations (1–2) equal to zero,

$$P^* = k_P\omega_{\rm sn}A \tag{6}$$

$$W^* = k_W\omega_{\rm sn}A. \tag{7}$$

Substituting the result for $P^*$ into equation (5), we get

$$\begin{aligned} C^* &= (1-\omega_{\rm sn})A\Big(Z_C + \frac{\alpha_C P^*/A}{\beta_C + P^*/A}\Big) \\ &= (1-\omega_{\rm sn})A\Big(Z_C + \frac{\alpha_C k_P\omega_{\rm sn}}{\beta_C + k_P\omega_{\rm sn}}\Big). \end{aligned} \tag{8}$$

#### Model equations – stochasticity

As in the case of continuous time, we use the assumption that the stochastic perturbations are small and independent between time steps. This allows us to apply the linear approximation. The dynamics for the deviations $\tilde{P}(t) = P(t) - P^*$, $\tilde{W}(t) = W(t) - W^*$ and $C(t) = C(t) - C^*$ are

$$\tilde{P}(t+1) = \tilde{P}(t) - r_P\tilde{P}(t) + \sigma_P^e u_P^e(t)P^* + \sigma_P^d u_P^d(t)\sqrt{P^*} \tag{9}$$

$$\tilde{W}(t+1) = \tilde{W}(t) - r_W\tilde{W}(t) + \sigma_W^e u_W^e(t)W^* + \sigma_W^d u_W^d(t)\sqrt{W^*} \tag{10}$$

$$\tilde{C}(t) = a_{CP}\tilde{P}(t) + \sigma_C^e u_C^e(t)C^*, \tag{11}$$

where

$$a_{CP} = (1-\omega_{\rm sn})\frac{\alpha_C\beta_C}{(\beta_C + P^*/A)^2}.$$

#### Variability equations

Under the assumption that the stochastic perturbations are small, the variance of pollinator biomass $P$ and wild plant biomass $W$ predicted by equations (9–10) is

$$\mathrm{Var}(P) = \frac{1}{r_P(2-r_P)}\Big(\big(\sigma_P^e P^*\big)^2 + \big(\sigma_P^d\sqrt{P^*}\big)^2\Big) \quad (12)$$

$$\mathrm{Var}(W) = \frac{1}{r_W(2-r_W)}\Big(\big(\sigma_W^e W^*\big)^2 + \big(\sigma_W^d\sqrt{W^*}\big)^2\Big), \quad (13)$$

so that the variability of pollinatior biomass $P$ and wild plant biomass $W$ is

$$\mathrm{CV}^2(P) = \frac{1}{r_P(2-r_P)}\Big((\sigma_P^e)^2 + \frac{(\sigma_P^d)^2}{P^*}\Big) \quad (14)$$

$$\mathrm{CV}^2(W) = \frac{1}{r_W(2-r_W)}\Big((\sigma_W^e)^2 + \frac{(\sigma_W^d)^2}{W^*}\Big). \quad (15)$$

The variation in crop yield has two independent contributions, see equation (11). Hence, the variance of crop yield is equal to the sum of the variances of these contributions,

$$\begin{aligned}\mathrm{Var}(C) &= a_{CP}^2\mathrm{Var}(P) + \big(\sigma_C^e C^*\big)^2 \\ &= \frac{1}{r_P(2-r_P)}\frac{\big((1-\omega_{\rm sn})\alpha_C\beta_C\big)^2}{(\beta_C+P^*/A)^4}\big((\sigma_P^e P^*)^2 + (\sigma_P^d)^2P^*\big) + \big(\sigma_C^e C^*\big)^2. \quad (16)\end{aligned}$$

Then, the variability of crop yield is

$$\begin{aligned}\mathrm{CV}^2(C) &= \frac{1}{r_P(2-r_P)}\frac{\big((1-\omega_{\rm sn})\alpha_C\beta_C\big)^2}{(\beta_C+k_P\omega_{\rm sn})^4}\frac{(\sigma_P^e P^*)^2 + (\sigma_P^d)^2P^*}{\big((1-\omega_{\rm sn})A\big(Z_C + \frac{\alpha_C k_P\omega_{\rm sn}}{\beta_C+k_P\omega_{\rm sn}}\big)\big)^2} + \big(\sigma_C^e\big)^2 \\ &= \frac{1}{r_P(2-r_P)}\frac{(\alpha_C\beta_C)^2}{(\beta_C+k_P\omega_{\rm sn})^4}\frac{\big(\sigma_P^e k_P\omega_{\rm sn}\big)^2 + (\sigma_P^d)^2 k_P\omega_{\rm sn}/A}{\big(Z_C + \frac{\alpha_C k_P\omega_{\rm sn}}{\beta_C+k_P\omega_{\rm sn}}\big)^2} + \big(\sigma_C^e\big)^2 \\ &= \frac{1}{r_P(2-r_P)}\frac{(\alpha_C\beta_C)^2}{(\beta_C+k_P\omega_{\rm sn})^2}\frac{\big(\sigma_P^e k_P\omega_{\rm sn}\big)^2 + (\sigma_P^d)^2 k_P\omega_{\rm sn}/A}{(Z_C\beta_C + (Z_C+\alpha_C)k_P\omega_{\rm sn})^2} + \big(\sigma_C^e\big)^2. \quad (17)\end{aligned}$$

It is instructive to consider the special case in which pollinator growth rate saturates, $r_P \approx c_P\alpha_P$, and demographic stochasticity of pollinators is negligible, $\sigma_P^d \approx 0$. Then,

$$\begin{aligned}\mathrm{CV}^2(C) &\approx \frac{1}{c_P\alpha_P(2-c_P\alpha_P)}\frac{(\alpha_C\beta_C)^2}{(\beta_C+k_P\omega_{\rm sn})^2}\frac{\big(\sigma_P^e k_P\omega_{\rm sn}\big)^2}{(Z_C\beta_C + (Z_C+\alpha_C)k_P\omega_{\rm sn})^2} + \big(\sigma_C^e\big)^2 \\ &= \frac{1}{c_P\alpha_P(2-c_P\alpha_P)}\frac{(\beta_C/k_P)^2}{(\beta_C/k_P+\omega_{\rm sn})^2}\frac{\big(\sigma_P^e\omega_{\rm sn}\big)^2}{(Z_C/\alpha_C\,\beta_C/k_P + (Z_C/\alpha_C+1)\omega_{\rm sn})^2} + \big(\sigma_C^e\big)^2 \\ &= \frac{\big(\sigma_P^e\big)^2}{c_P\alpha_P(2-c_P\alpha_P)}\left[\frac{\beta_C/k_P}{\beta_C/k_P+\omega_{\rm sn}}\frac{\omega_{\rm sn}}{Z_C/\alpha_C\,\beta_C/k_P + (Z_C/\alpha_C+1)\omega_{\rm sn}}\right]^2 + \big(\sigma_C^e\big)^2. \quad (18)\end{aligned}$$

This shows that the dependence of the variability of crop yield on the proportion of semi-natural habitat is effectively controlled by two parameter combinations: $Z_C/\alpha_C$ and $\beta_C/k_P$.

### Supplementary Methods 3. Connecting spatial and non-spatial models, and derivation of mean-field approximation

#### Non-spatial model

Deterministic dynamics,

$$P(t+1) = P(t) \exp\Big(r_P\big(1 - \frac{P(t)}{k_P \omega_{\mathrm{sn}} A}\big)\Big) \tag{1}$$

$$W(t+1) = W(t) \exp\Big(r_W\big(1 - \frac{W(t)}{k_W \omega_{\mathrm{sn}} A}\big)\Big), \tag{2}$$

with

$$r_P = \frac{\alpha_P \big(W(t) + C(t)\big)}{\beta_P + W(t) + C(t)} \tag{3}$$

$$r_W = \frac{\alpha_W P(t)/A}{\beta_W + P(t)/A}. \tag{4}$$

Crop yield is instantaneous function,

$$C(t) = (1 - \omega_{\mathrm{sn}}) A \Big(Z_C + \frac{\alpha_C P(t)/A}{\beta_C + P(t)/A}\Big). \tag{5}$$

#### Spatial model

Deterministic dynamics for semi-natural habitat cell,

$$P_k(t+1) = P_k(t) \exp\Big(r_{P,k}\big(1 - \frac{P_k(t)}{k_P A_1}\big)\Big) \tag{6}$$

$$W_k(t+1) = W_k(t) \exp\Big(r_{W,k}\big(1 - \frac{W_k(t)}{k_W A_1}\big)\Big), \tag{7}$$

with $A_1$ the area of a single cell and

$$r_{P,k} = \frac{\alpha_P \big(\sum_{\ell \notin L} \varepsilon_{k\ell} W_\ell(t) + \sum_{\ell \in L} \varepsilon_{k\ell} C_\ell(t)\big)}{\beta_P + \sum_{\ell \notin L} \varepsilon_{k\ell} W_\ell(t) + \sum_{\ell \in L} \varepsilon_{k\ell} C_\ell(t)} \tag{8}$$

$$r_{W,k} = \frac{\alpha_W \sum_{\ell \notin L} \varepsilon_{k\ell} P_\ell(t)/A}{\beta_W + \sum_{\ell \notin L} \varepsilon_{k\ell} P_\ell(t)/A}, \tag{9}$$

with $L$ the set of crop land cells. Crop yield is instantaneous function,

$$C_k(t) = A_1 \Big(Z_C + \frac{\alpha_C \sum_{\ell \notin L} \varepsilon_{k\ell} P_\ell(t)/A}{\beta_C + \sum_{\ell \notin L} \varepsilon_{k\ell} P_\ell(t)/A}\Big). \tag{10}$$

The distance-decay function is

$$\varepsilon_{k\ell} = \frac{1 + \beta}{1 + \beta \exp(\alpha d k\ell)}, \tag{11}$$

with $dk\ell$ the distance between cells $k$ and $\ell$, and parameters $\alpha$ and $\beta$ determining the 50% decay distance $dm$ and the 90% to 10% decay distance $\Delta d$ (see main text and Appendix 1).

#### Without distance-decay

Take $\alpha \to 0$, then $\varepsilon_{k\ell} = 1$ and

$$r_{P,k} = \frac{\alpha_P\big(\sum_{\ell\notin L} W_\ell(t) + \sum_{\ell\in L} C_\ell(t)\big)}{\beta_P + \sum_{\ell\notin L} W_\ell(t) + \sum_{\ell\in L} C_\ell(t)} = \frac{\alpha_P\big(W(t)+C(t)\big)}{\beta_P + W(t) + C(t)} = r_P \tag{12}$$

$$r_{W,k} = \frac{\alpha_W \sum_{\ell\notin L} P_\ell(t)/A}{\beta_W + \sum_{\ell\notin L} P_\ell(t)/A} = \frac{\alpha_W P(t)/A}{\beta_W + P(t)/A} = r_W \tag{13}$$

$$C_k(t) = A_1\Big(Z_C + \frac{\alpha_C \sum_{\ell\notin L} P_\ell(t)/A}{\beta_C + \sum_{\ell\notin L} P_\ell(t)/A}\Big) = A_1\Big(Z_C + \frac{\alpha_C P(t)/A}{\beta_C + P(t)/A}\Big). \tag{14}$$

These equations imply that the spatial model without distance-decay reduces exactly to the non-spatial model (at least the deterministic part).

#### With distance-decay

In equilibrium,

$$P_k = k_P A_1 = P_1 \quad \text{and} \quad P = k_P\, \omega_{\text{sn}} A \tag{15}$$

$$W_k = k_W A_1 = W_1 \quad \text{and} \quad W = k_W\, \omega_{\text{sn}} A \tag{16}$$

and

$$C_k = A_1 Z_C + A_1 \frac{\alpha_C \sum_{\ell\notin L} \varepsilon_{k\ell} P_1/A}{\beta_C + \sum_{\ell\notin L} \varepsilon_{k\ell} P_1/A} \tag{17}$$

$$\begin{aligned} C &= (1-\omega_{\text{sn}})AZ_C + A_1\alpha_C \sum_{k\in L} \frac{\sum_{\ell\notin L} \varepsilon_{k\ell}}{\beta_C A/P_1 + \sum_{\ell\notin L}\varepsilon_{k\ell}} \\ &= (1-\omega_{\text{sn}})AZ_C + \frac{A\alpha_C}{n^2} \sum_{k\in L} \frac{\frac{1}{n^2}\sum_{\ell\notin L}\varepsilon_{k\ell}}{\beta_C/k_P + \frac{1}{n^2}\sum_{\ell\notin L}\varepsilon_{k\ell}}. \end{aligned} \tag{18}$$

The formula indicates that the effects of the spatial fragmentation pattern on crop yield are modulated by two factors: the ratio $\beta_C/k_P$ and the distance-decay $\varepsilon_{k\ell}$.

Consider the sums $\sum_{\ell\notin L}\varepsilon_{k\ell}$. There are $\omega_{\text{sn}} n^2$ terms (possible values of $\ell$), and $(1-\omega_{\text{sn}})n^2$ such sums (possible values of $k$). Define

$$\overline{\varepsilon} = \frac{1}{(1-\omega_{\text{sn}})n^2}\frac{1}{\omega_{\text{sn}} n^2} \sum_{k\in L, \ell\notin L} \varepsilon_{k\ell}, \tag{19}$$

i.e., the average value of $\varepsilon_{k\ell}$ when taking a random cell $k \in L$ and a random cell $\ell \notin L$. Then the "mean-field" approximation (physics jargon) is

$$\begin{aligned} C &\approx (1-\omega_{\text{sn}})AZ_C + (1-\omega_{\text{sn}})A\alpha_C \frac{\omega_{\text{sn}}\overline{\varepsilon}}{\beta_C/k_P + \omega_{\text{sn}}\overline{\varepsilon}} \\ &= (1-\omega_{\text{sn}})A\Big(Z_C + \frac{\alpha_C\omega_{\text{sn}}\overline{\varepsilon}}{\beta_C/k_P + \omega_{\text{sn}}\overline{\varepsilon}}\Big). \end{aligned}$$

Comparing this with the solution of the non-spatial model

$$C = (1-\omega_{\text{sn}})A\Big(Z_C + \frac{\alpha_C\omega_{\text{sn}}}{\beta_C/k_P + \omega_{\text{sn}}}\Big), \tag{20}$$

we see that the spatial model structure effectively increases the parameter $\beta_C/k_P$ to $\beta_C/(\overline{\varepsilon}k_P)$ (recall that $\overline{\varepsilon} \le 1$). As expected, we recover the non-spatial model result by setting $\overline{\varepsilon} = 1$.

#### Interpretation of $\bar{\varepsilon}$

Multiply $\bar{\varepsilon}$ by the area of semi-natural habitat, $R_P = \omega_{\mathrm{sn}} A \,\bar{\varepsilon} = \omega_{\mathrm{sn}} n^2 A_1 \,\bar{\varepsilon}$, then $R_P$ is the area from which a crop plant can be pollinated, averaged over all crop plants (i.e. averaged over all crop land cells).

Multiply $\bar{\varepsilon}$ by the crop land area, $R_C = (1-\omega_{\mathrm{sn}}) A \,\bar{\varepsilon} = (1-\omega_{\mathrm{sn}}) n^2 A_1 \,\bar{\varepsilon}$, then $R_C$ is the crop land area that a pollinator can reach, averaged over all pollinators (i.e. averaged over all semi-natural habitat cells).

Independently of the interpretation, the mathematical anaylsis shows that the effective parameter $\beta_C/k_P$ of the non-spatial model has to be replaced by

$$\frac{1}{\bar{\varepsilon}} \frac{\beta_C}{k_P} = \frac{\omega_{\mathrm{sn}} A}{R_P} \frac{\beta_C}{k_P} = \frac{(1-\omega_{\mathrm{sn}}) A}{R_C} \frac{\beta_C}{k_P}$$

to effectively take into account the spatial structure.

Some special cases ($\sqrt{A}$ and $\sqrt{A_1}$ are the linear dimensions of the landscape and of a cell, respectively):

- For $d_m > \sqrt{A}$, we have $\bar{\varepsilon} \to 1$ and we recover the non-spatial model.
- For $d_m < \sqrt{A_1}$, we have $\bar{\varepsilon} \to 0$, and there is no pollination.
- For $d_m \sim \sqrt{A_1}$, only crop plants close to a fragment boundary can be pollinated. Then $R_P \approx d_m^2$ for crop plants close to a fragment boundary and $R_P = 0$ otherwise. Hence, averaged over all crop plants, $R_P \approx \frac{b\, d_m}{(1-\omega_{\mathrm{sn}}) A} d_m^2$ and $\bar{\varepsilon} \approx \frac{b\, d_m^3}{(1-\omega_{\mathrm{sn}})\, \omega_{\mathrm{sn}} A^2}$ with $b$ the total perimeter of all fragments. Hence, for fixed $\omega_{\mathrm{sn}}$, $\bar{\varepsilon}$ is determined by the fragments perimeter.
- For "random" fragmentation patterns, the fraction of semi-natural habitat cells in any subset of the landscape is equal to $\omega_{\mathrm{sn}}$. Hence, for all crop plants, $R_P \approx \omega_{\mathrm{sn}} d_m^2$ and $\bar{\varepsilon} \approx \frac{d_m^2}{A}$. Note that $\bar{\varepsilon}$ is independent of $\omega_{\mathrm{sn}}$ in this case.

#### Some illustrations

We consider three types of fragmentation pattern: "dispersed" (first row), "random" (second row) and "clustered" (third row). Semi-natural habitat in green, crop land in yellow.

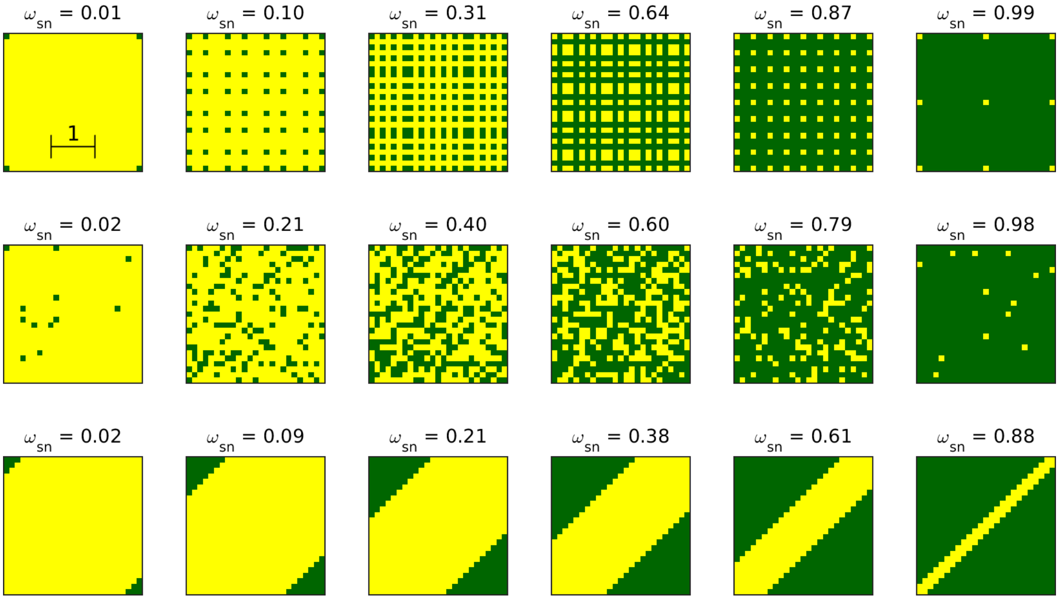

Note the scale in the top-left panel, to be compared with the values of $d_m$ below.

We plot pollination-dependent crop yield (crop pollination for short) as a function of proportion of semi-natural habitat $\omega_{\rm sn}$. Rows: different values of $d_m = \Delta d$. Columns: different values of $\beta_C/k_P$. Each panel shows results for the three types of fragmentation pattern: circles for dispersed, squares for random and $\times$-marks for clustered.

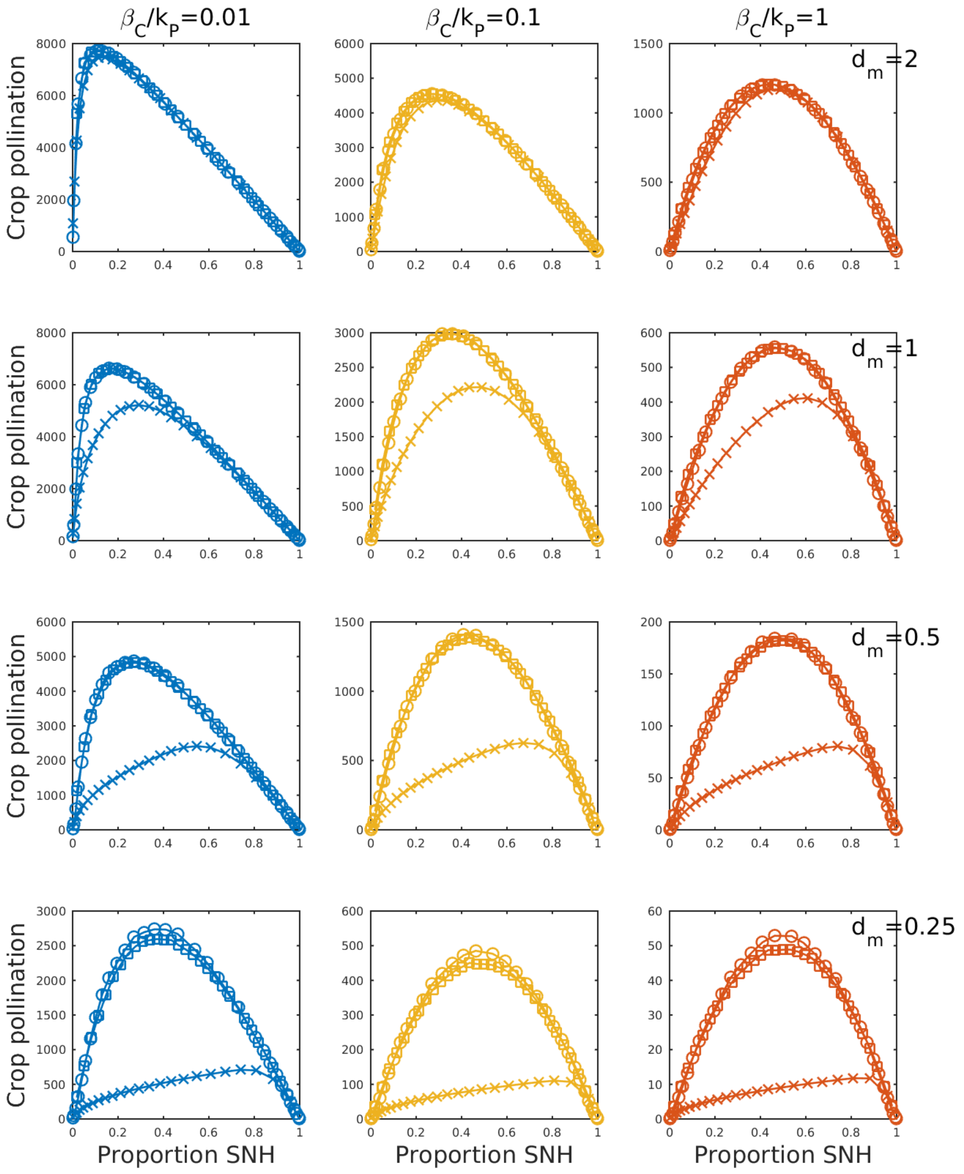

For large $d_m$, the results for the three pattern types coincide, and tend towards those of the non-spatial model. For smaller $d_m$, the results for the dispersed and random pattern type are close, while those for the clustered pattern type are different (smaller). There are quantitative differences with respect to the main text; the reason being that kP does not depend on fragmentation (it is a fixed parameter here). This, however, does not affect the general fact that the mean-field approximation is a very accurate description of the model dynamics and its exact solution (Figure 2, main text) .

We have a closer look at the mean-field approximation. For $\beta_C/k_P = 0.1$, we plot crop pollination as a function of proportion of semi-natural habitat $\omega_{\text{sn}}$. Rows: different values of $d_m = \Delta d$. Columns: different pattern types: dispersed in first column, random in second column, and clustered in third column. Full line: exact solution; symbols (circles, squares, $\times$-marks): mean-field approximation.

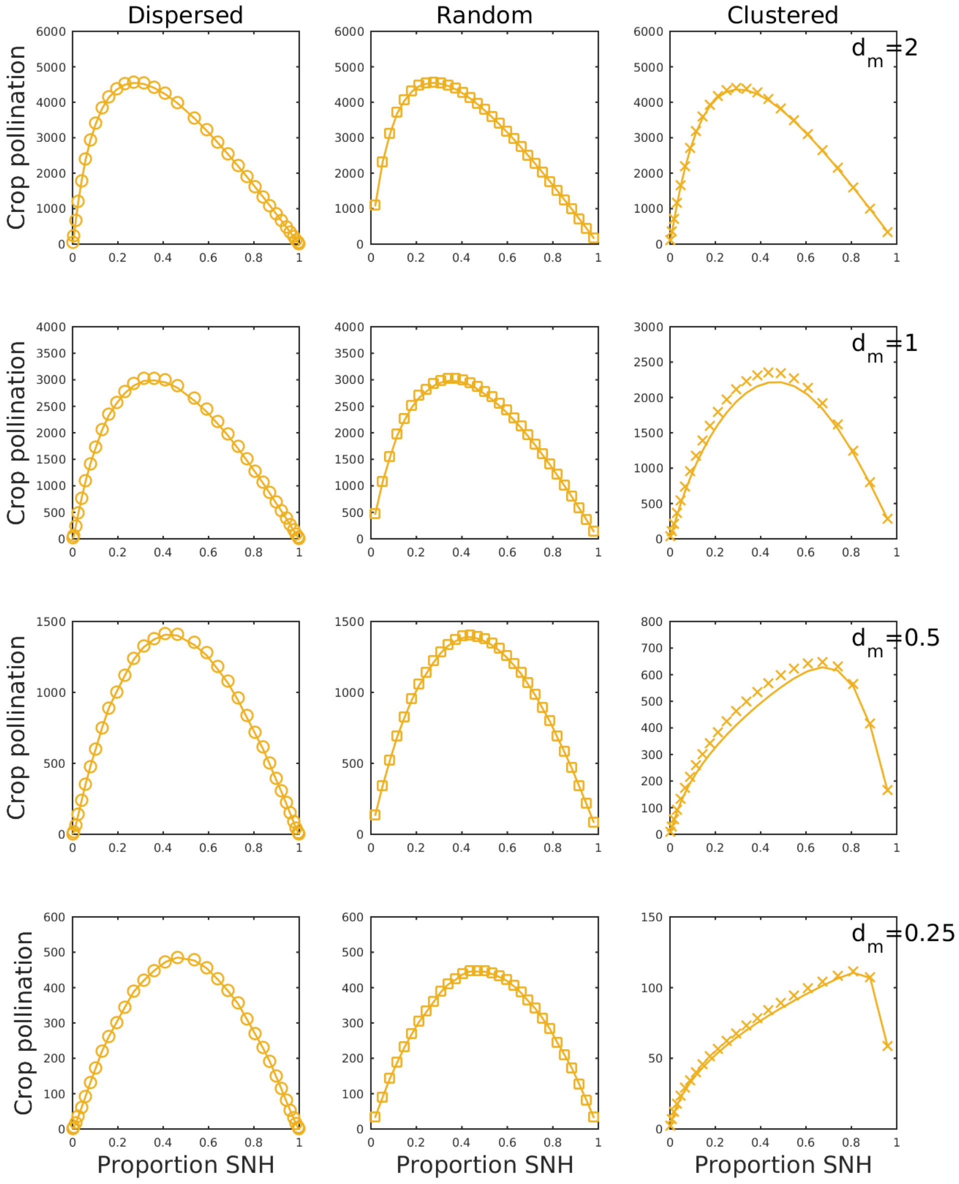

The accuracy of the mean-field approximation is very good; only for the clustered patterns there are visible differences. This suggests that in this model the spatial structure can be taken into account by modifying the parameter $\beta_C/k_P$, i.e., by increasing it to $\beta_C/(\bar{\varepsilon}k_P)$.

Hence, the problem of understanding the effects of fragmentation splits up into two sub-problems. First, we have to understand the relationship between fragmentation pattern and effective distance-decay $\bar{\varepsilon}$. Second, we have to understand the relationship between effective distance-decay $\bar{\varepsilon}$ and crop pollination. Both questions are addressed in the main text.

### Supplementary Table and Figures

**Table S1**
**Figures S1-S15**

**Table S1.** Parameters and variables of the model

| Parameters & Variables | Definition | Dimensions |
|---|---|---|
| Parameters | | |
| $\alpha_P$ | Maximum growth rate of pollinators | $time^{-1}$ |
| $\alpha_W$ | Maximum growth rate of semi-natural plants | $time^{-1}$ |
| $\alpha_C$ | Maximum crop yield derived from pollinator interactions | $mass \cdot area^{-1}$ |
| $\beta_P$ | Half-saturation constant of pollinators | mass |
| $\beta_W$ | Half-saturation constant of 'wild' plants | $mass \cdot area^{-1}$ |
| $\beta_C$ | Half-saturation constant of crop plants to pollinators | $mass \cdot area^{-1}$ |
| $k_P$ | Carrying capacity of pollinators per unit area | $mass \cdot area^{-1}$ |
| $k_W$ | Carrying capacity of semi-natural plants per unit area | $mass \cdot area^{-1}$ |
| $A$ | Total landscape area | area |
| $A_1$ | Area of a single cell | area |
| $\omega_{sn}$ | Proportion of semi-natural habitat | dimensionless |
| $Z_C$ | Crop yield independent of pollinators | $mass \cdot area^{-1}$ |
| $r_{P,k}$ (t) | Intrinsic growth rate of pollinators in cell $k$ | $time^{-1}$ |
| $r_{W,k}$ (t) | Intrinsic growth rate of 'wild' plants in cell $k$ | $time^{-1}$ |
| $r_{C,k}$ (t) | Intrinsic growth rate of crops in cell $k$ | $time^{-1}$ |
| $\varepsilon_{kl}$ | Distance decay function of ecosystem service flow | distance |
| $d_{kl}$ | Distance between cells $k$ and $l$ | distance |
| $d_m$ | Distance from seminatural cell edge at which the pollination flow equals one half of its initial value | distance |
| $\Delta d$ | Distance over which the pollination flow decreases from 90% to 10% of its initial value | distance |
| $\alpha$, $\beta$ | Auxiliary parameters to determine $d_m$ and $\Delta d$ | dimensionless |
| $b$ | Parameter modulating the effect of the metapopulation capacity. It reflects the ability of species to live in fragmented landscapes | dimensionless |
| $\lambda_M$ | Metapopulation capacity of the fragmented landscape | $mass \cdot area^{-1}$ |
| $c_s$ | Parameter of the SAR function | $(mass \cdot individuals)/area^2$ |
| $z_s$ | Parameter of the SAR function | dimensionless |
| $c_{kp}$ | Parameters of the power law ($k_P$ dependence on $S$) | $(mass \cdot area^{-1})$/ species |
| $z_{kp}$ | Parameters of the power law ($k_P$ dependence on $S$) | dimensionless |
| $\sigma^e_P$ | Environmental standard deviation of pollinators | $time^{-1/2}$ |
| $\sigma^e_W$ | Environmental standard deviation of 'wild' plants | $time^{-1/2}$ |
| $\sigma^e_C$ | Environmental standard deviation of crop production | dimensionless |

| | | |
|---|---|---|
| $\sigma^d_P$ | Demographic standard deviation of pollinators | $mass^{1/2}\cdot time^{-1/2}$ |
| $\sigma^d_W$ | Demographic standard deviation of semi-natural plants | $mass^{1/2}\cdot time^{-1/2}$ |
| $u^e_P$, $u^d_P$,<br>$u^e_W$, $u^d_W$,<br>$u^e_C$, $u^d_C$ | White noise signals with zero mean and standardized variance. $u^e$ = environmental, $u^d$ = demographic<br>$P$ = pollinators; $W$ = 'wild' plants; $C$ = crop plants | dimensionless |
| Variables | | |
| $C_k\ (t)$ | Biomass of crop plants (crop yield) in cell $k$ | mass |
| $W_k\ (t)$ | Biomass of semi-natural or 'wild' plants in cell $k$ | mass |
| $P_k\ (t)$ | Biomass of pollinators in cell $k$ | mass |
| $S$ | Number of pollinator species | # individuals |

**Figure S1**. Effects of landscape composition and MPP on ecosystem services. Results correspond to exponential distance-decay of pollination flow. Ecosystem services are represented as a function of the proportion of seminatural habitat, for different MPP. MPP includes the effects of fragmentation – more specifically, the aggregation pattern of land conversion – and the distance-decay of ecosystem service flow. Parameter values: $\alpha_P = \alpha_W = 0.9$, $\beta_P = \beta_W = 0.6$, $A = 10$, $Z_C = 1000$, $\alpha_C = 1000$, $k_W = 5000$, $k_P = 0.1$, $e_P = 0.8$, $\sigma^d_P = 0.1$, $\sigma^e_C = 0.03$, $\alpha_C = 1000$, Pollination dependence = 50%, $z_{k_P} = 0.26$.

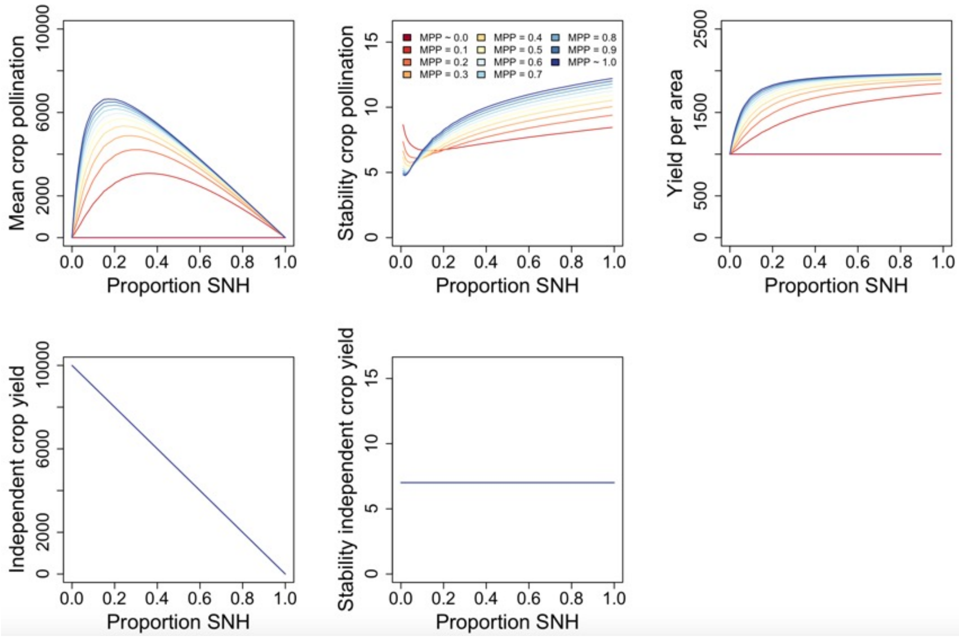

**Figure S2**. Effects of $\Delta d$ on ecosystem services. Pollination dependence = 50%; $d_m = 1$. Top row plots correspond to random land conversion patterns ($w$=0). Bottom row correspond to aggregated land conversion patterns ($w$=5).

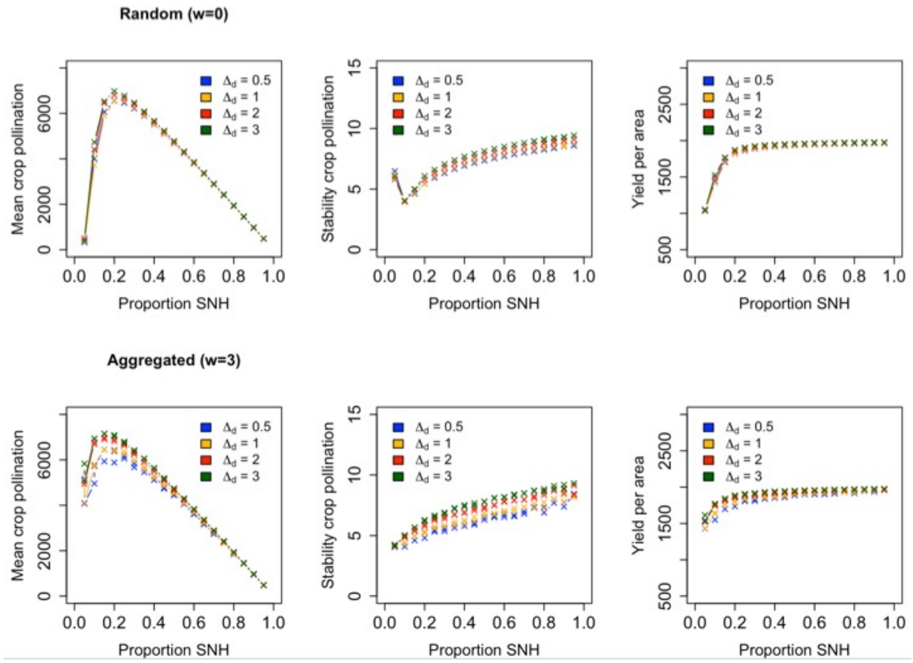

**Figure S3**. Example of land conversion pattern generation. Variation in the values of *w*, *m* allowed us to produce a continuous gradient of land conversion patterns, and therefore fragmentation, based on the level of aggregation (see main text). All figures represent different fragmentation scenarios for the same proportion of seminatural habitat (50%): green cells correspond to seminatural habitat, whereas yellow cells represent crop land.

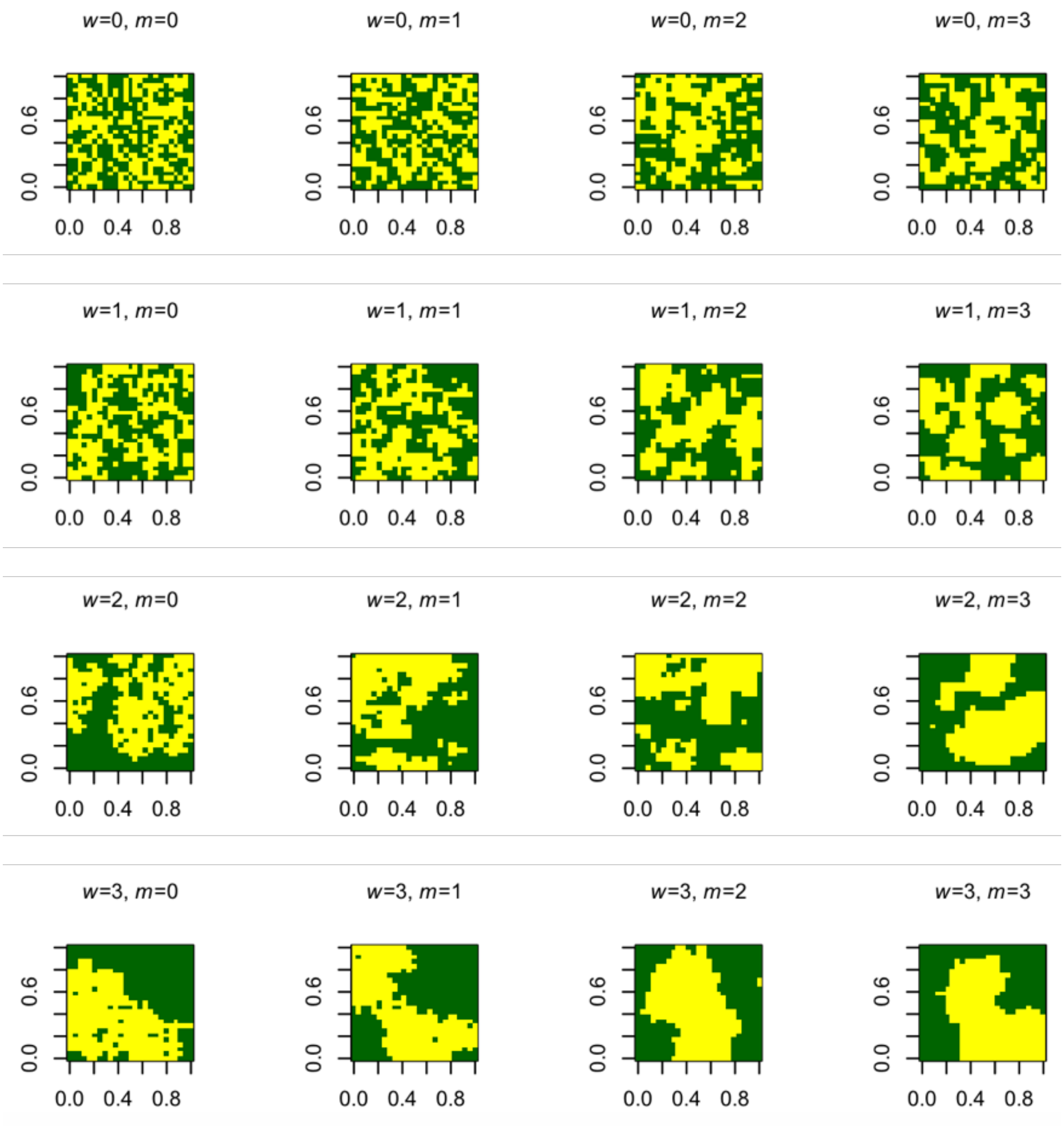

**Figure S4**. Mean Pollination Potential (MPP) and crop's relative requirement for pollinators ($\beta_C/k_P$), and their effects on crop pollination. When $\beta_C/k_P$ is small, crop yield saturates at lower pollinator biomass than their carrying capacity; when $\beta_C/k_P$ is large, crop yield saturates at pollinator biomasses much higher than their carrying capacities. (**A**) Effects of $\beta_C/k_P$ on the mean and stability of crop pollination. On one hand, greater values of $\beta_C/k_P$ increase the effect of pollinator biomass on crop pollination, reducing mean yield and shifting maximum yield to larger amounts of seminatural habitat. On the other hand, $\beta_C/k_P$ controls how fast the saturation of crop pollination to pollinator biomass sets in and, consequently, how fast the response of crops to pollinator stochasticity drops down; thus, the smaller $\beta_C/k_P$ the faster the saturation sets in, and so the faster crop yield variability drops when increasing seminatural habitat (**B**) Effects of varying MPP on $\beta_C/k_P$ and the mean and stability of crop pollination. Without distance-decay (or when MPP ≈ 1), the spatial model collapses into the non-spatial model. Fragmentation effects on ecosystem services become stronger when MPP<1, which increases $\beta_C/k_P$.

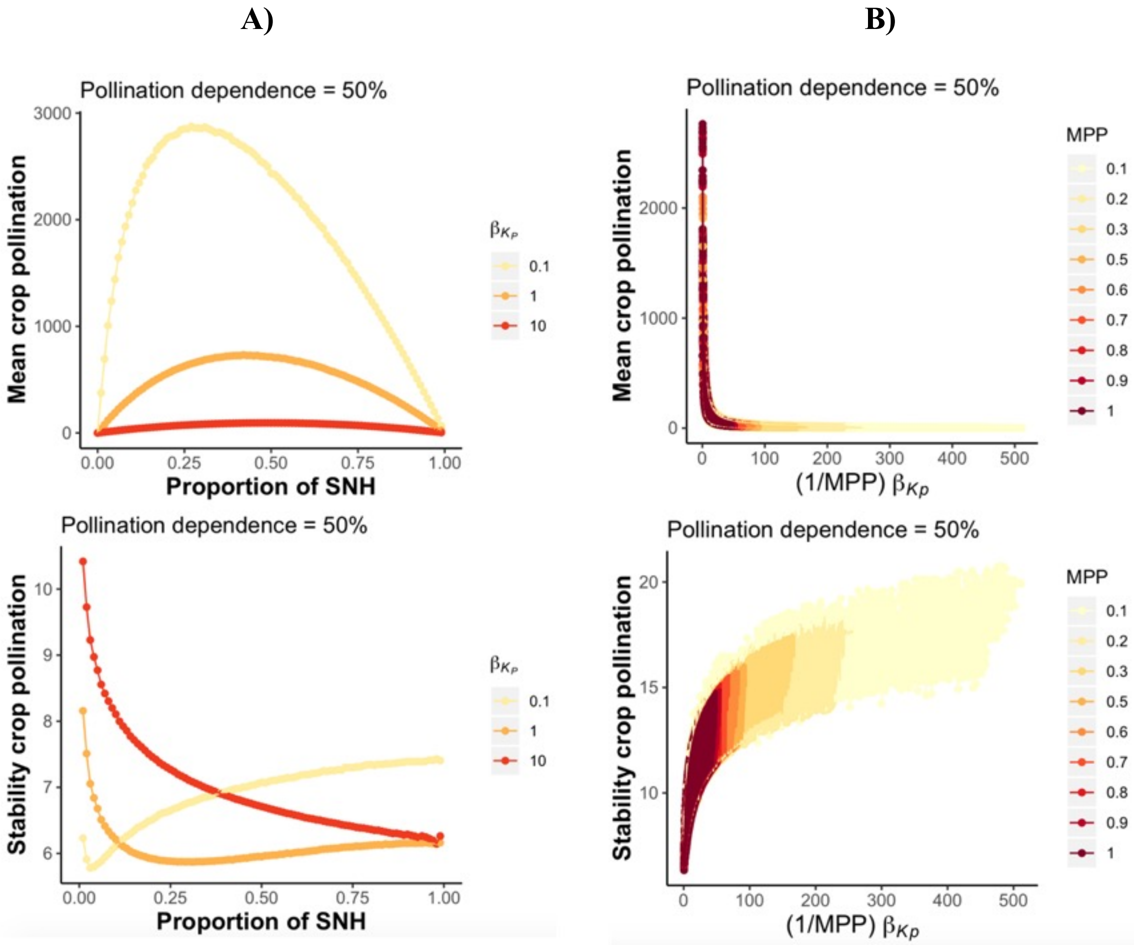

**Figure S5**. Effects of fragmentation on MPP. In general, more fragmented (less aggregated) areas of seminatural habitat result in higher MPP, especially with fast distance-decay of service flows (low $d_m$).

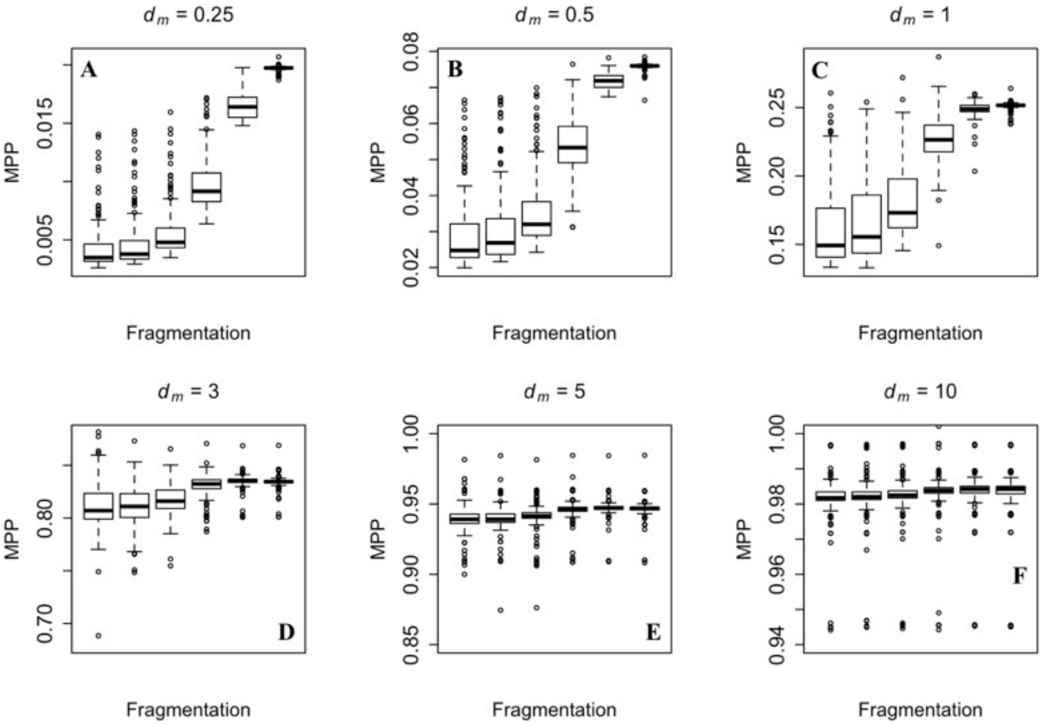

**Figure S6**. Effects of distance-decay ($d_m$) on MPP

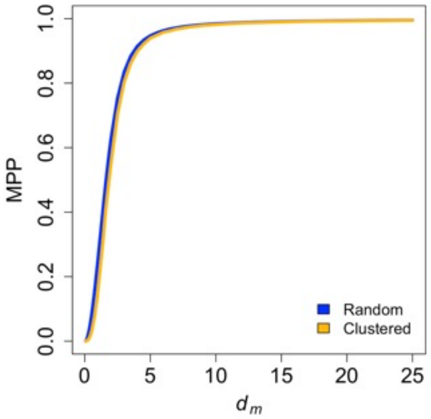

**Figure S7**. Effect of fragmentation metrics on ecosystem services. The effects of fragmentation metrics on several services associated with crop production are not clear. **A)** Random land conversion (more fragmented). **B)** Aggregated land conversion

**A)**

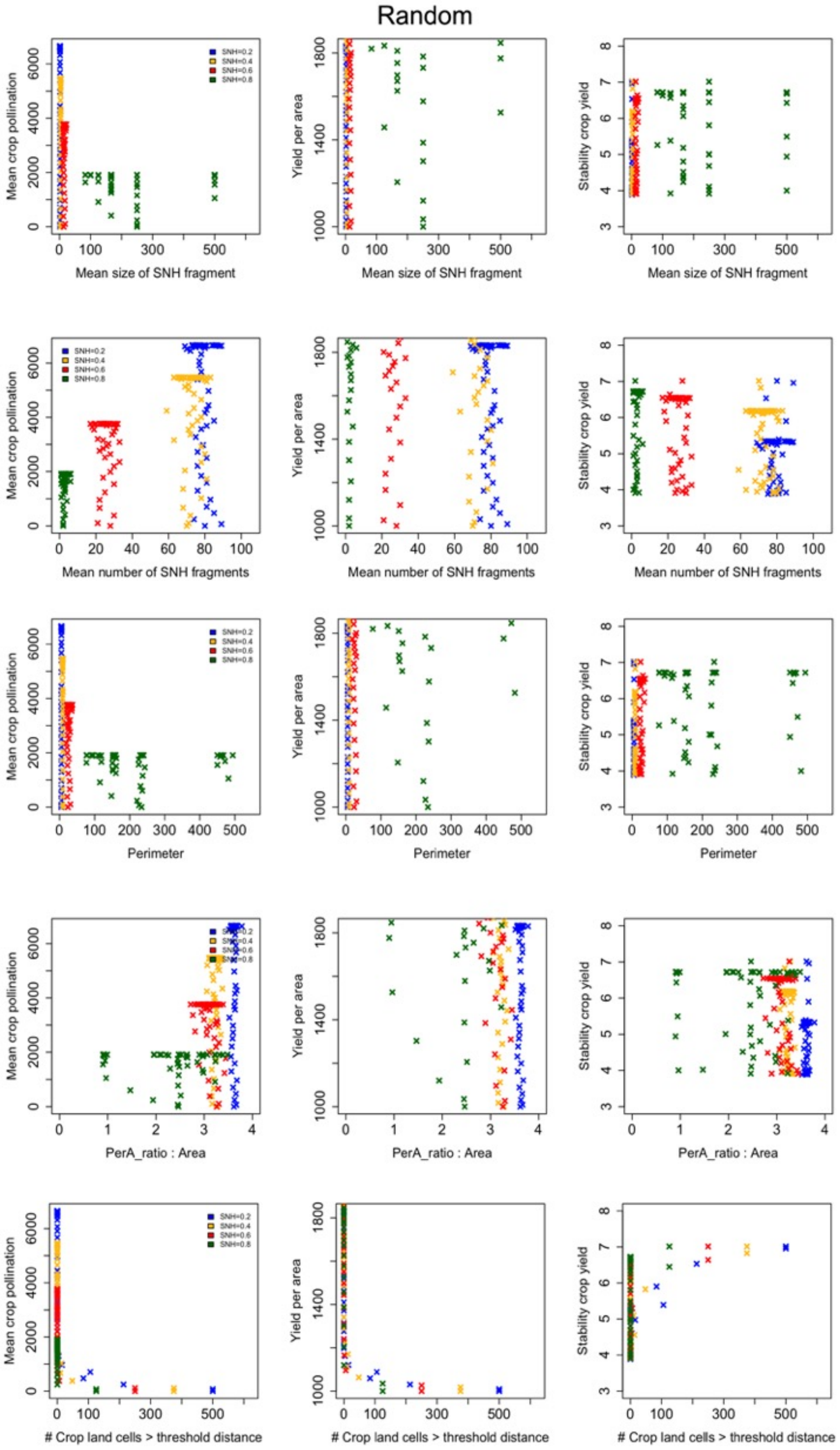

**B)**

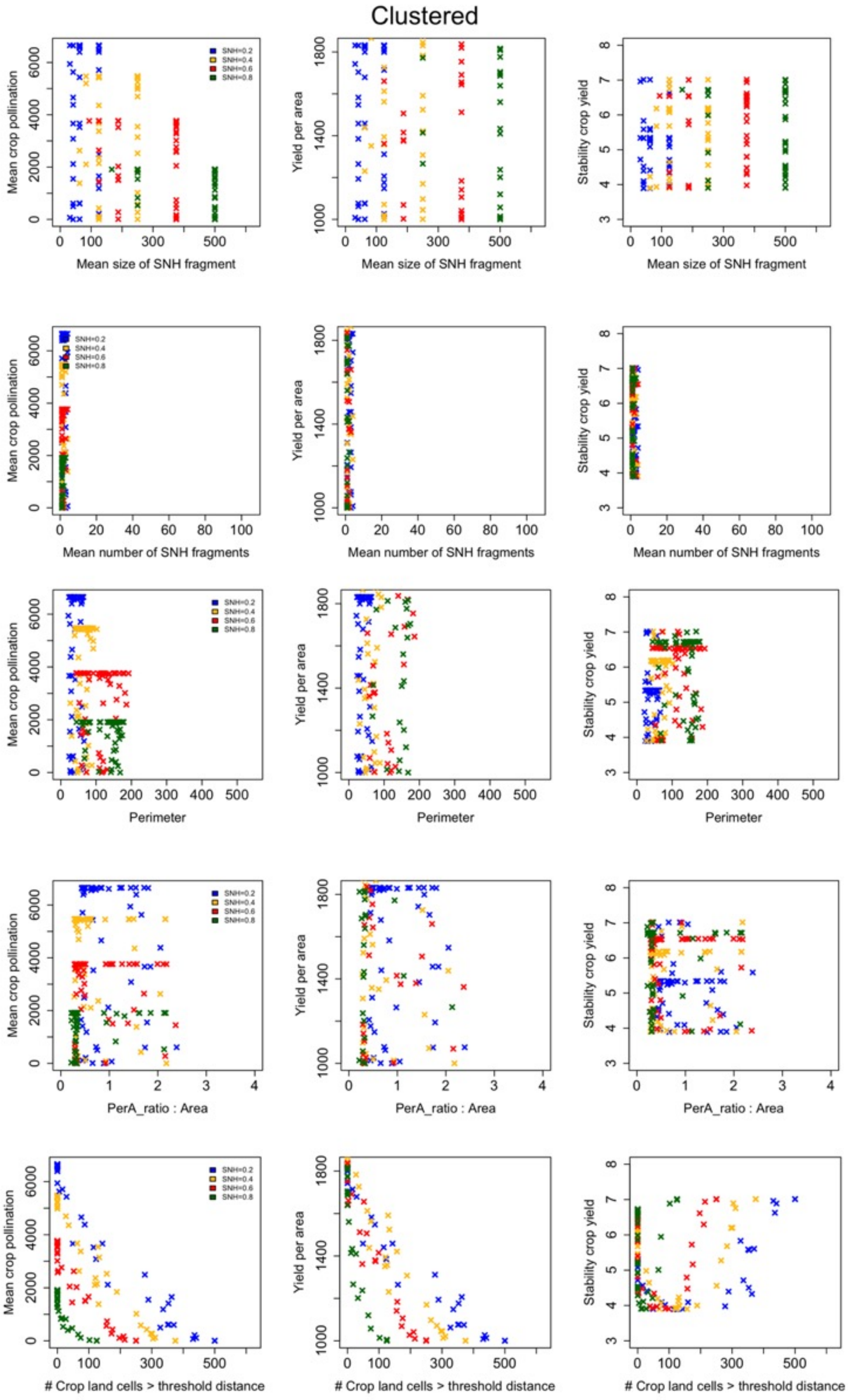
Clustered
SNH=0.2
SNH=0.4
SNH=0.6
SNH=0.8
Mean crop pollination
Yield per area
Stability crop yield
Mean size of SNH fragment
Mean number of SNH fragments
Perimeter
PerA_ratio : Area
### Crop land cells > threshold distance

**Figure S8**. Effect of Mean Pollinator Potential (MPP) on ecosystem services. **A)** Random land conversion. **B)** Aggregated land conversion.

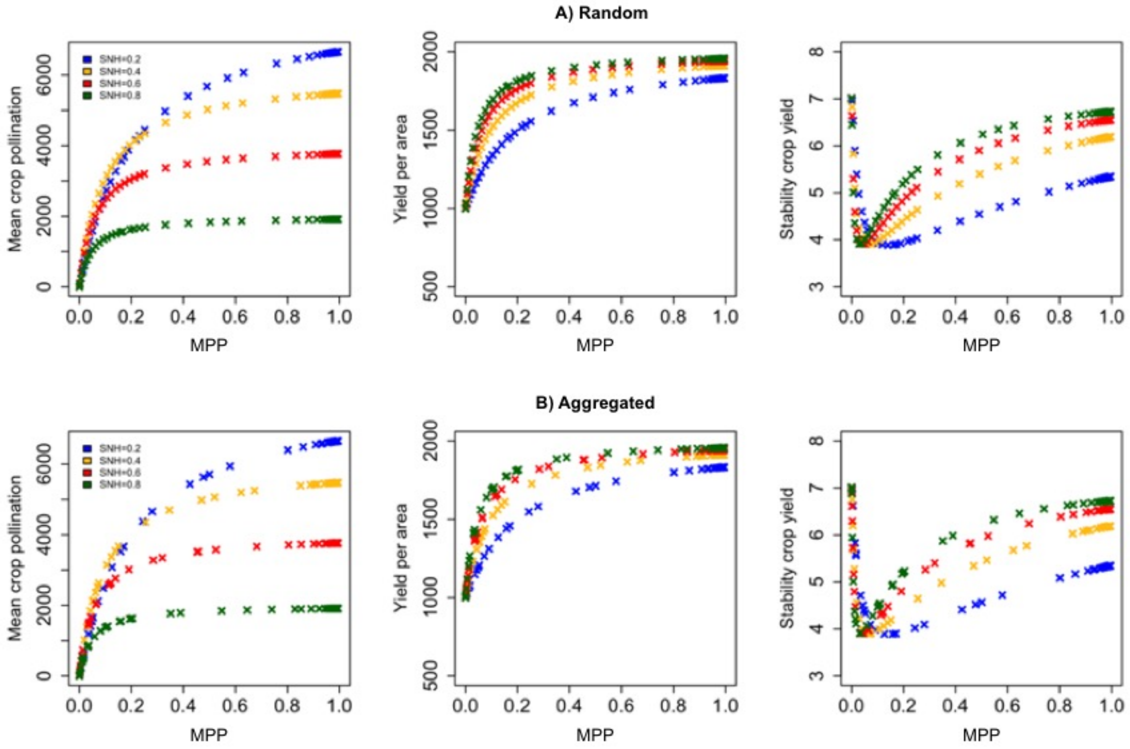

**Figure S9**. Effects of landscape composition and MPP on ecosystem services. Pollination dependence = 50%. **A)** $z_{k_P}$= 0; **B)** $z_{k_P}$= 0.5. A higher biodiversity effect (larger $z_{k_P}$) increases both mean crop pollination and its stability, as well as yield per area. Ecosystem services are represented as a function of the proportion of seminatural habitat, for different MPP. MPP includes the effects of fragmentation – more specifically, the aggregation pattern of land conversion – and the distance-decay of ecosystem service flow. Parameter values: $\alpha_P = \alpha_W = 0.9$, $\beta_P = \beta_W = 0.6$, $A = 10$, $Z_C = 1000$, $\alpha_C = 1000$, $k_W = 5000$, $e_P = 0.8$, $\sigma^d_P = 0.1$, $\sigma^e_C = 0.03$, $\alpha_C = 1000$, Pollination dependence = 50%.

**A)**

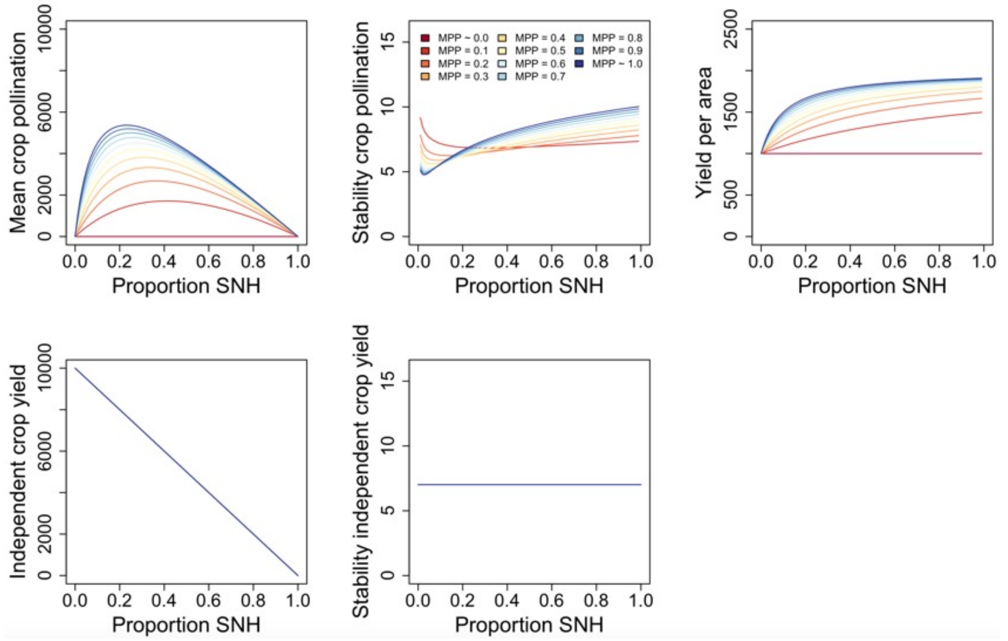

**B)**

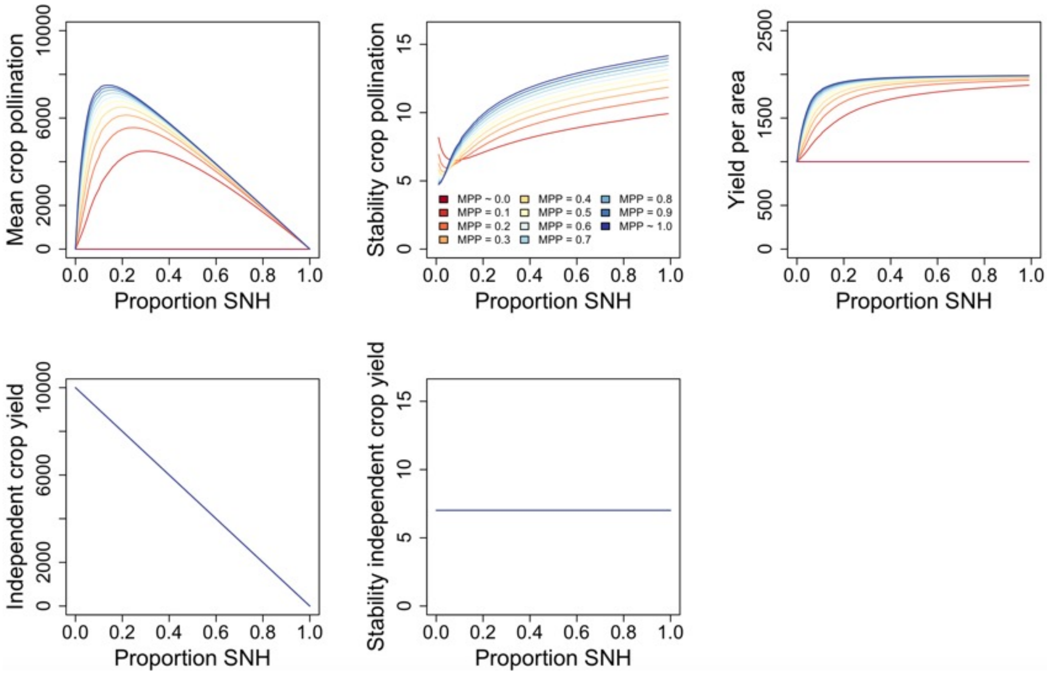

**Figure S10**. Effects of crop pollination dependence on ecosystem services. **A**) Pollination dependence = 10%. **B**) Pollination dependence = 90%. Ecosystem services are represented as a function of the proportion of seminatural habitat, for different MPP. MPP includes the effects of fragmentation – more specifically, the aggregation pattern of land conversion – and the distance-decay of ecosystem service flow. Parameter values: $\alpha_P = \alpha_W = 0.9$, $\beta_P = \beta_W = 0.6$, $A = 10$, $k_W = 5000$, $e_P = 0.8$, $\sigma^d{}_P = 0.1$, $\sigma^e{}_C = 0.03$, $\alpha_C = 1000$, $z_{k_P} = 0.26$.

**A) Pollination dependence = 10%**

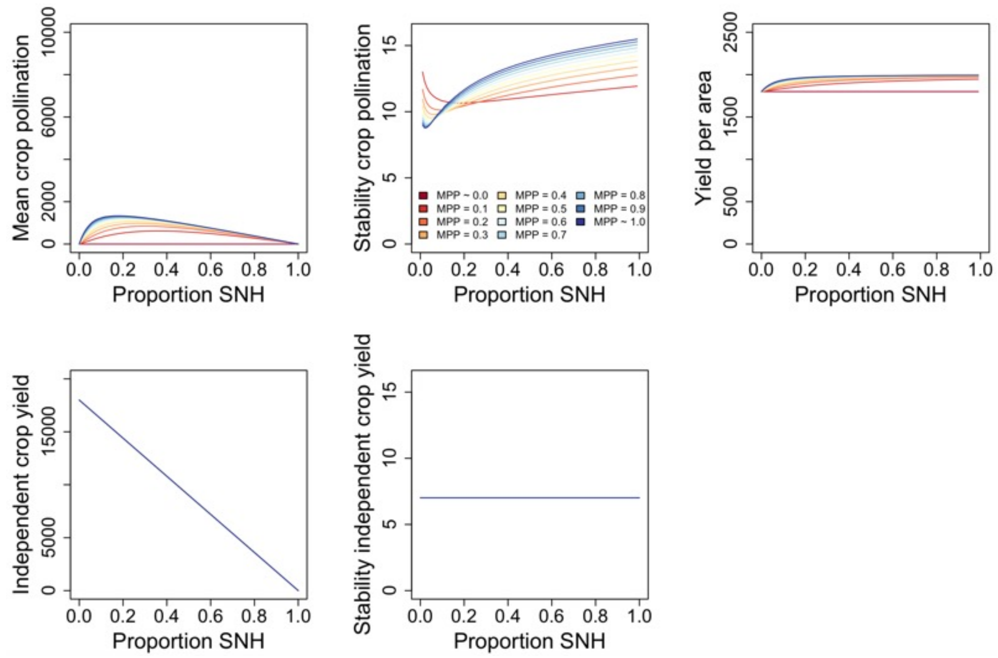

**A) Pollination dependence = 90%**

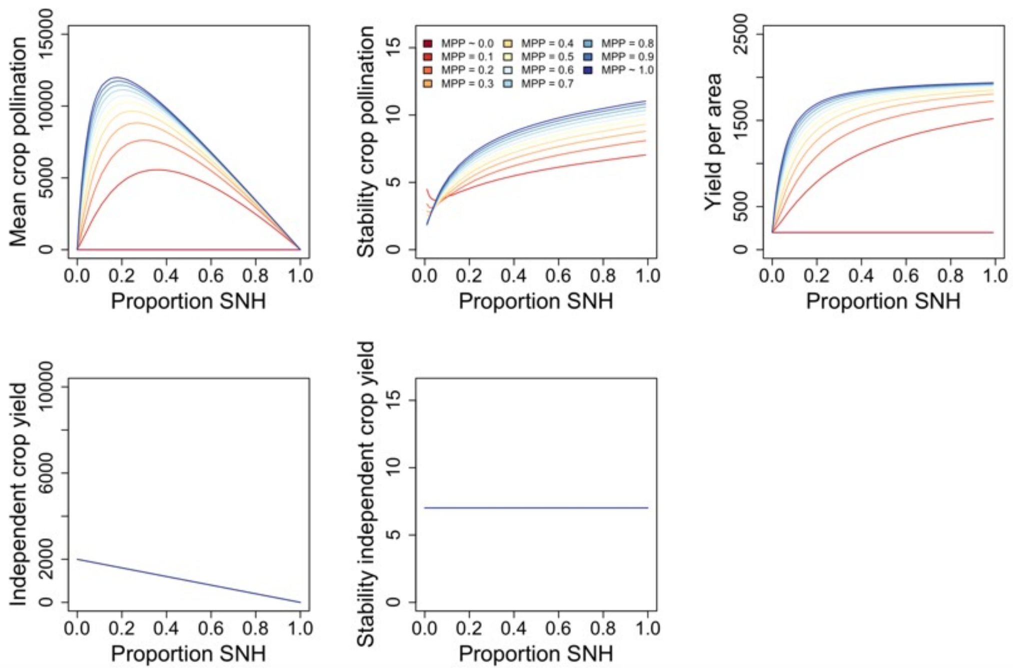

**Figure S11**. Effects of aggregation on biodiversity. $d_m = 1$, $\Delta d = 1$, dispersal distance = 1, $b = 10$. For simplicity, we set $w_a = w_b$ and include only $w$ in the legend.

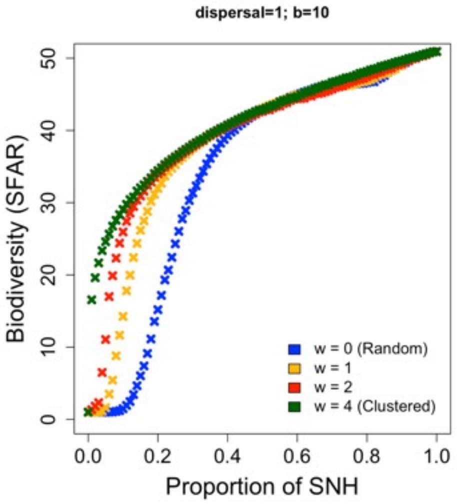

**Figure S12**. Effects of dispersal distance and $b$ on biodiversity. $d_m = 1$, $\Delta d = 1$. For simplicity, we set $w_a = w_b$ and include only $w$ in the legend.

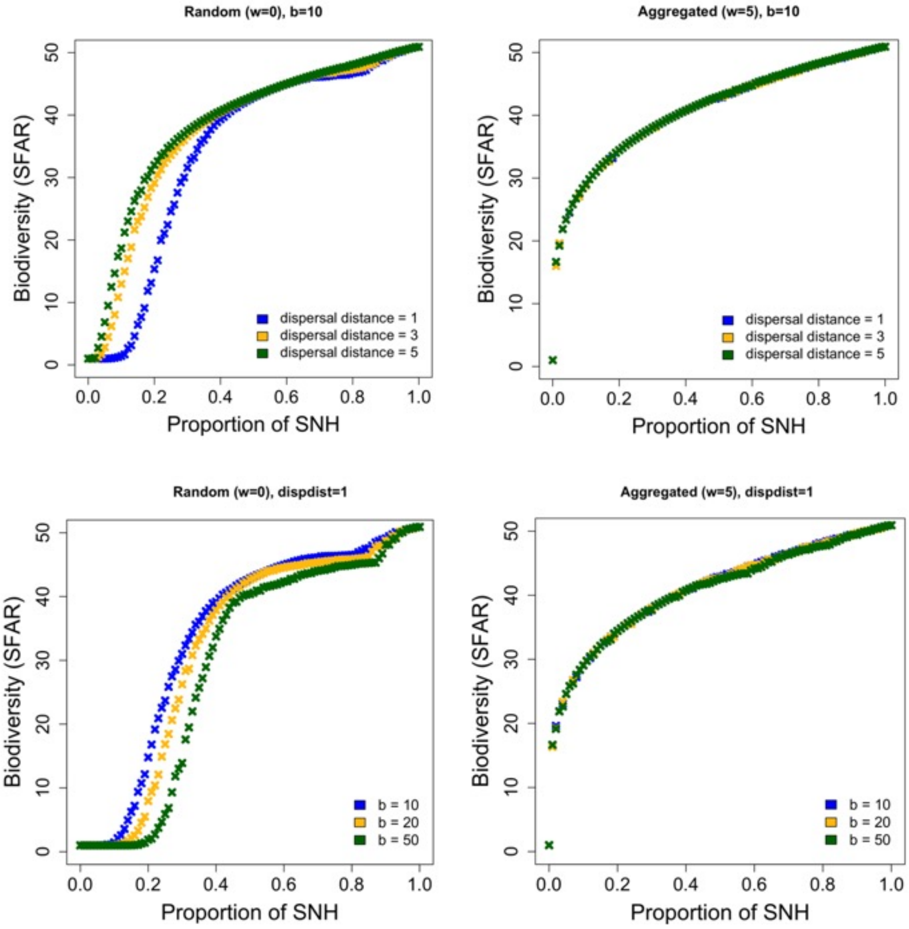

**Figure S13**. Effects of changing $h$ on crop pollination. Plots show the response of crop pollination services – mean and stability of crop pollination (subpanels A and B), and yield per area (subpanel C) – as a function of the proportion of seminatural habitat (SNH). All MPP values are contained within the shadows, whose limits are determined by the minimum and maximum values across the range of MPP. Biodiversity can affect crop pollination services by increasing the carrying capacity of pollinators ($k_p = c_{k_P} S^{z_{k_P}}$) and reducing the response of crop production to environmental fluctuations ($\sigma_P^e = e_P / S^q$). Light orange and grey shadows reflect, respectively, the response of ecosystem services in a scenario where environmental stochasticity depends on biodiversity ($q = 1/2$; Tilman 1999) with another scenario where biodiversity does not affect environmental stochasticity ($q = 0$). Each panel represent two cases based on the magnitude of the effect of biodiversity on the pollinator's carrying capacity. In panel **A**, this effect is null ($z_{k_P}$= 0), whereas in panel **B** biodiversity has a large effect on crop pollination ($z_{k_P}$= 0.5). Parameter values: $\alpha_P = \alpha_W = 0.9$, $\beta_P = \beta_W = 0.6$, $A = 10$, $Z_C = 1000$, $\alpha_C = 1000$, $k_W = 5000$, $e_P = 0.8$, $\sigma^d{}_P = 0.1$, $\sigma^e{}_C = 0.03$, $\alpha_C = 1000$, Pollination dependence = 50%.

**A)**

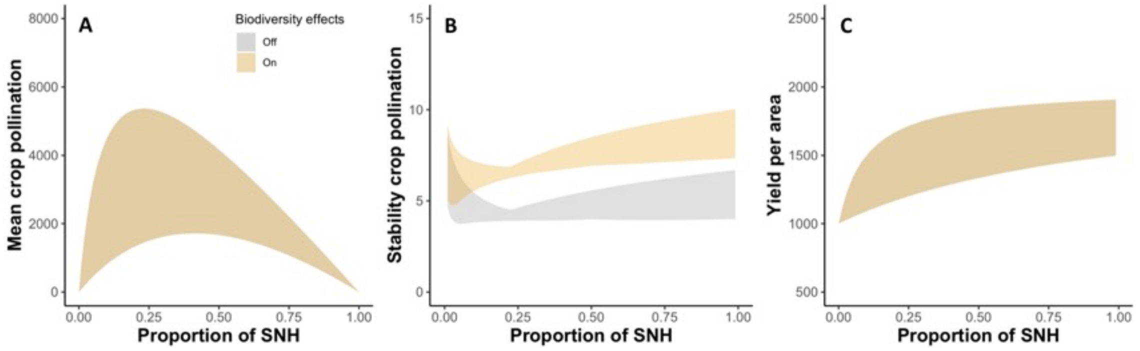

**B)**

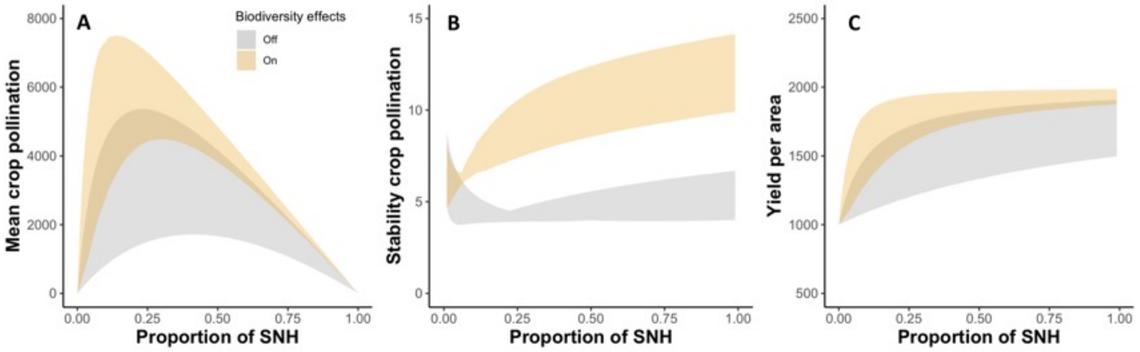

**Figure S14**. Effects of fragmentation (aggregation of seminatural habitat fragments) on pollinator-dependent ecosystem services. Ecosystem services are plotted as a function of the proportion of seminatural habitat for different fragmentation levels. For simplicity, we set $w = m$ (Random land conversion: $w = m = 0$; Contiguous land conversion: $w, m > 0$; higher $w$, $m$ means more aggregation). Rows represent increasing values of decay distance $d_m$ (0.5, 1, 5). Parameter values: $\alpha_P = \alpha_W = 0.9$, $\beta_P = \beta_W = 0.6$, $A = 10$, $Z_C = 1000$, $\alpha_C = 1000$, $k_W = 5000$, $e_P = 0.8$, $\sigma^d{}_P = 0.1$, $\sigma^e{}_C = 0.03$, $\alpha_C = 1000$, Pollination dependence = 50%, $z_{k_P} = 0.26$.

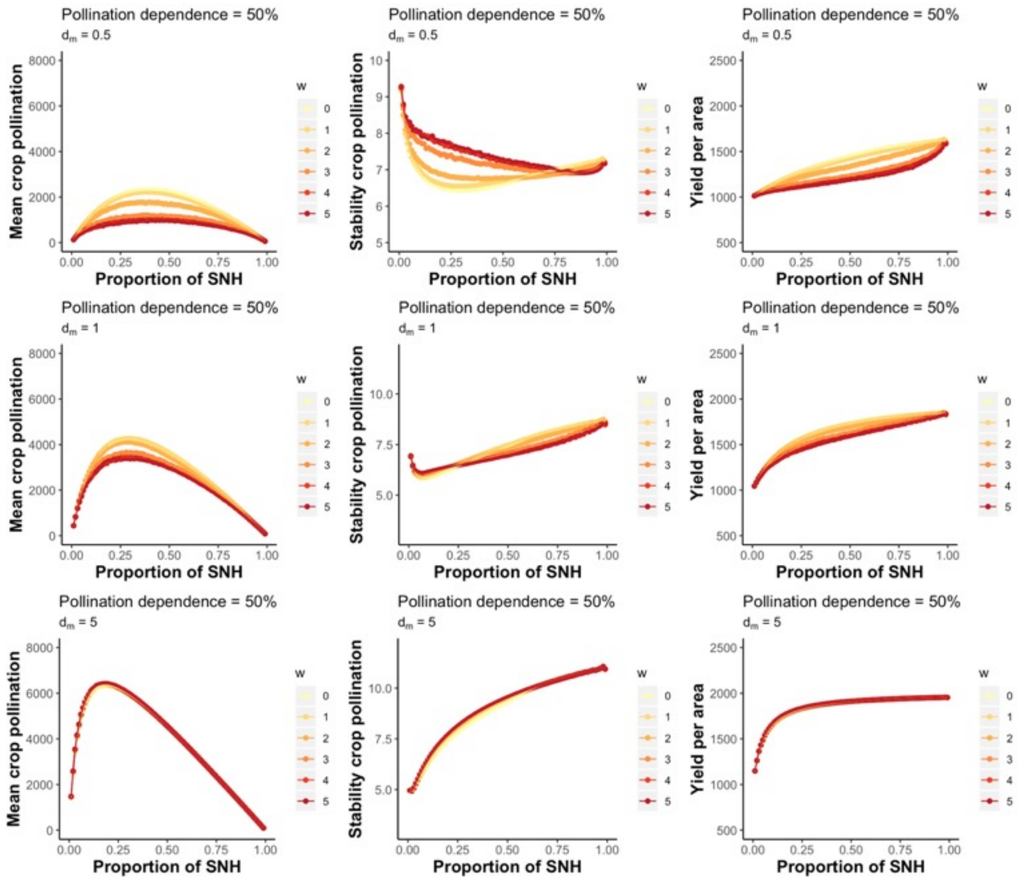

**Figure S15**. Net effects of aggregation on crop pollination services. Ecosystem services are plotted as a function of fragmentation for different proportion of seminatural habitat or SNH (as opposed to figure 3). Pollination dependence = 50%. Rows represent increasing values of decay distance $d_m$ (0.5, 1, 5). **A)** $z_{k_P}$= 0; **B)** $z_{k_P}$= 0.5. Parameter values: $\alpha_P = \alpha_W = 0.9$, $\beta_P = \beta_W = 0.6$, $A = 10$, $Z_C = 1000$, $\alpha_C = 1000$, $k_W = 5000$, $e_P = 0.8$, $\sigma^d_P = 0.1$, $\sigma^e_C = 0.03$, $\alpha_C = 1000$, Pollination dependence = 50%.

**A)**

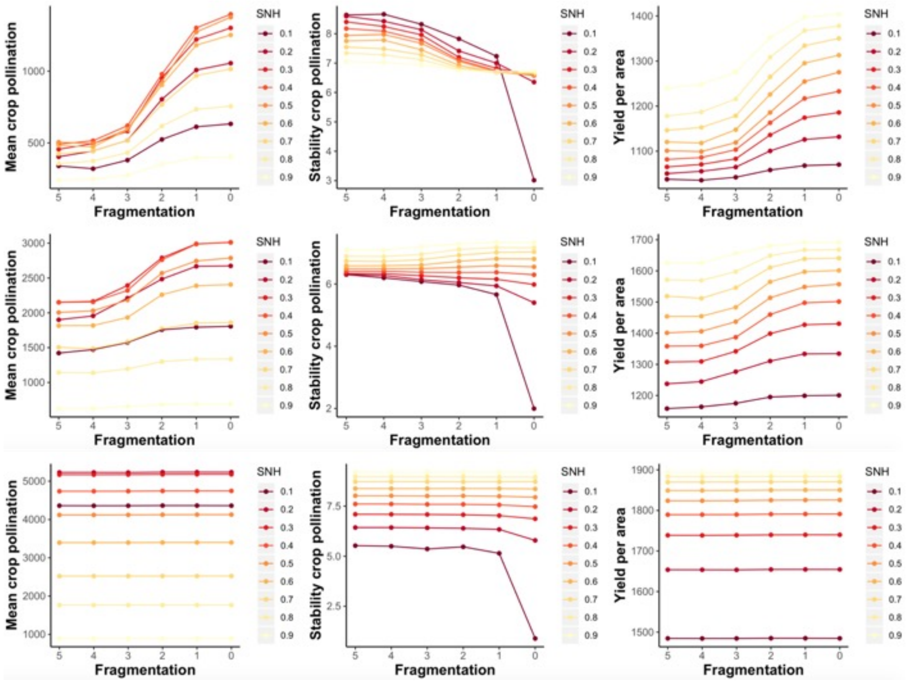

**B)**

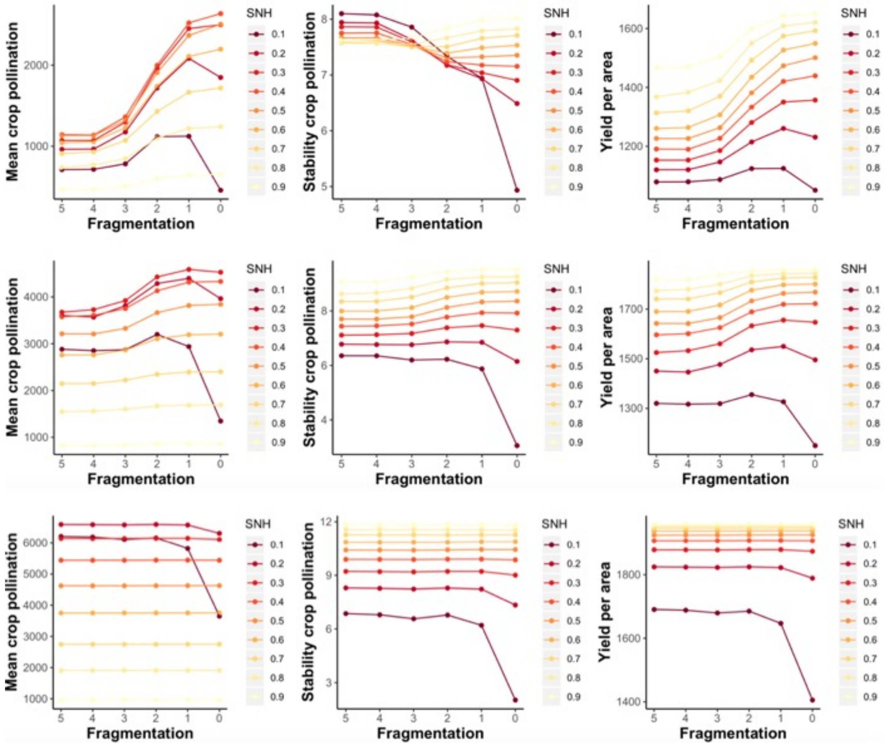
